# The mycomembrane proteins PorH and ProtX are inserted at polar growth zones and linked to the cell wall

**DOI:** 10.1101/2025.10.14.682376

**Authors:** Elizabeth M. Hart, Dieuwertje A. de Bruin, Victoria M. Marando, Marina Arnau Alemany, Eric D. Snow, Bailey J. Schultz, Erkin Kuru, Suzanne Walker, Andrea Vettiger, Thomas G. Bernhardt

**Author notes:** Corresponding authors: Thomas G. Bernhardt, Andrea Vettiger, Elizabeth M. Hart. These authors contributed equally to this work (Elizabeth M. Hart, Dieuwertje A. de Bruin). Artificial intelligence disclosure: AlphaFold2 was used in the research for this manuscript.

## Abstract

The Mycobacteriales order of bacteria includes important pathogens such as *Mycobacterium tuberculosis*. These organisms are surrounded by a unique cell envelope architecture that includes a two-layered cell wall composed of peptidoglycan (PG) and arabinogalactan (AG). They also build an outer membrane called the mycomembrane that is made of mycolic acids. Mycolate outer membrane proteins (MOMPs) reside within the mycomembrane and a subset are thought to form pores that allow essential nutrients to permeate the envelope. However, little is known about the structure of these proteins or the mechanism by which they are assembled. Here, we investigate MOMP assembly in *Corynebacterium glutamicum* (*Cglu*) using PorH as a model corynebacterial MOMP. PorH is encoded in an operon with the MOMP PorA, and the two small, alpha-helical proteins have been proposed to form hetero-oligomeric pores in the mycomembrane. Consistent with this proposal, AlphaFold2 predicts a high confidence structure of a hetero-oligomeric pore formed by five copies each of PorH and its partner PorA, and we show that PorA is required for the surface assembly of PorH. Using a fluorescence assay for detection of surface-exposed PorH or another MOMP called ProtX, we found that MOMP assembly occurs within zones of active PG synthesis at the cell poles. We also discovered that PorH and ProtX are linked to the cell wall. Thus, like Gram-negative bacteria, *Cglu* coordinates outer membrane protein assembly with PG biogenesis and uses proteins to connect the mycomembrane and the cell wall.

**SIGNIFICANCE:** Diderm bacteria in the Mycobacteriales order have a distinctive outer layer called the mycomembrane. Proteins that reside within the mycomembrane play critical roles in virulence and cell viability. However, how proteins are assembled into the mycomembrane has remained an outstanding question in the field. Here, we investigate the biogenesis of mycomembrane proteins in the model organism *Corynebacterium glutamicum*. We show that these proteins are inserted into the mycomembrane at sites of nascent peptidoglycan growth and are attached to the cell wall. Thus, mycomembrane protein assembly shares surprisingly similar features with that of outer membrane protein assembly in Gram-negative bacteria like *Escherichia coli* despite their distant evolutionary relationship and the distinct properties of the proteins and membranes involved.

## INTRODUCTION

The Mycobacteriales order of bacteria includes pathogens such as *Mycobacterium tuberculosis*, *Mycobacterium abscessus*, and *Corynebacterium diphtheriae*, as well as environmental bacteria like *Corynebacterium glutamicum* and *Mycobacterium smegmatis*. Bacteria in the Mycobacteriales order assemble an elaborate cell envelope architecture with a second membrane made of mycolic acid lipids that distinguishes them from other microbes in the Actinomycetota phylum. These organisms are therefore referred to non-taxonomically as the mycolata and their corresponding envelope architecture as the mycolata envelope. Like most bacteria, the foundation of the envelope is a plasma membrane surrounded by a cell wall made of peptidoglycan (PG). Among the Mycobacteriales, the PG layer is decorated with a second glycan polymer called the arabinogalactan (AG) to form a composite wall structure. Mycolic acids are covalently attached to the AG component of the cell wall. Together with trehalose-linked mycolic acids and other diffusible lipids, these AG-bound mycolic acids form the second membrane structure called the mycomembrane (MM) (**Fig. 1A**) (1, 2). This multi-layered and interconnected envelope serves both as a target for antibiotics and as a formidable barrier preventing drug penetration (3). Detailed characterization of the pathways responsible for assembly of the mycolata envelope therefore has the potential to reveal new ways of targeting mycobacterial and corynebacterial pathogens with antibiotics.

**Figure 1:**
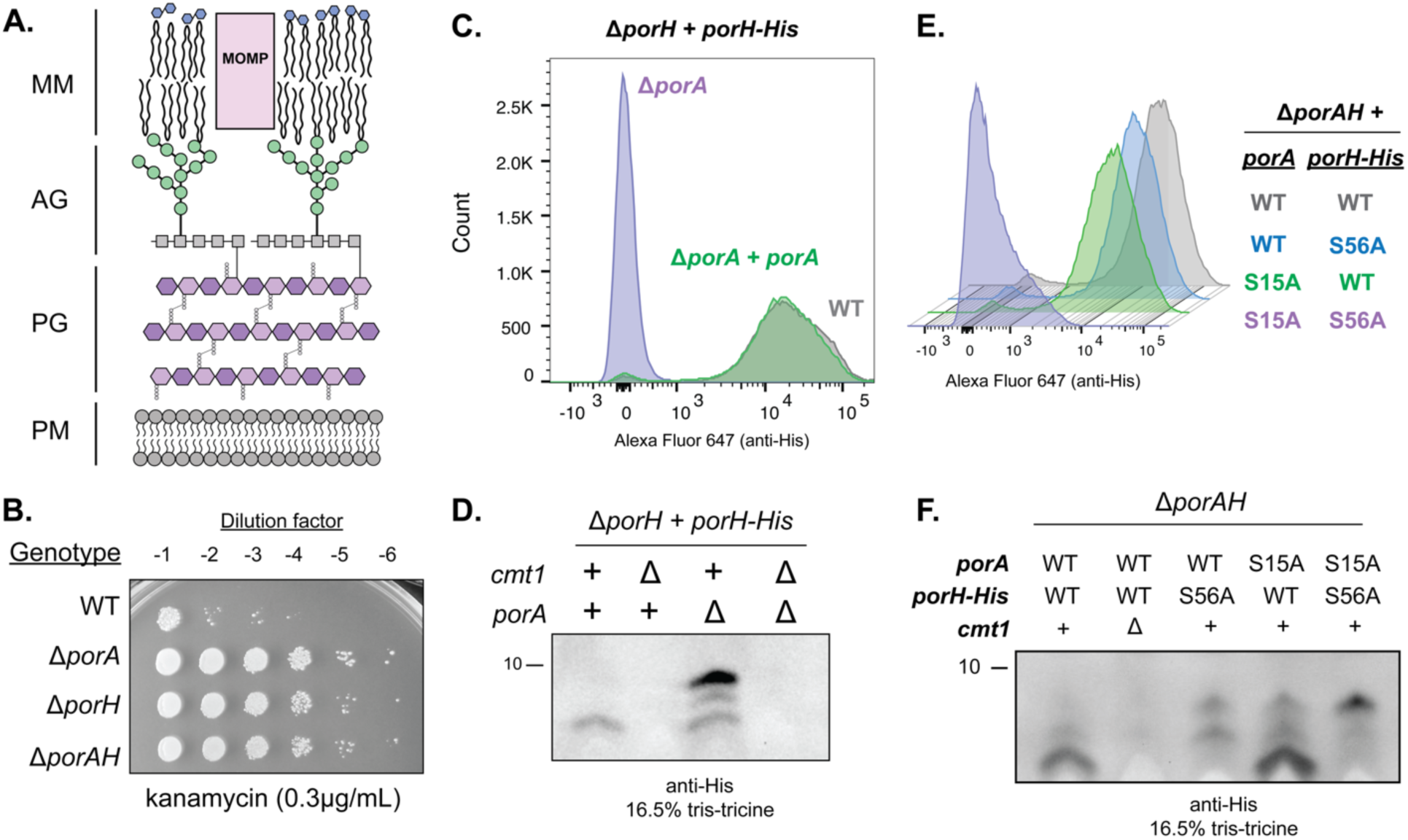
PorA and lipidation of either PorA or PorH are required for PorH surface assembly. **(A)** Depiction of the mycolata cell envelope. MM: mycomembrane, MOMP: mycolate outer membrane protein, AG: arabinogalactan, PG: peptidoglycan, PM: plasma membrane. **(B-F)** All experiments were performed in biological triplicate and a representative replicate is displayed. **(B)** Ten-fold serial dilutions of the indicated cultures were spotted onto brain heart infusion (BHI) medium supplemented with 9.1% sorbitol containing 0.3μg/mL kanamycin. **(C)** Flow cytometry analysis of PorH-His surface exposure in the indicated genetic backgrounds. Cells were grown to mid-log and stained with anti-6x His antibody labeled with Alexa Fluor 647. A representative replicate is shown as a histogram. All strains have the native copy of *porH* deleted and are complemented with constitutively expressed *porH-His* on a replicating vector. Complementation of Δ*porA* was performed through constitutive ectopic expression of *porA* from the same replicating vector and promoter as *porH-His*. **(D)** Immunoblot analysis of PorH-His in the indicated strains. The “+” sign indicates a wild-type allele and the “Δ” sign indicates a deletion/disruption. In all strains, the native copy of *porH* was deleted and constitutively complemented *in trans* from a replicating vector. **(E)** Flow cytometry of the indicated strains lacking the native copies of *porAH* and constitutively expressing *porH-His/porA* alleles from a replicating vector. Cells were grown to mid-log and stained with anti-6x His antibody labeled with Alexa Fluor 647 antibody. A representative replicate is shown as a histogram **(F)** Immunoblot analysis of the indicated strains, which are the same as those analyzed in **(C).** The “+” or “WT” label indicates a wild-type allele, the “Δ” symbol indicates a deletion of the indicated gene, and “S56A, S15A, or S56A/S15A” annotations indicate the mutated *O*-mycoloylation residue(s).

Despite the importance of the mycolata cell envelope in cell survival and pathogenicity, many aspects of its biogenesis remain unclear. One major gap in our understanding involves the mechanisms responsible for the transport and assembly of proteins within the MM. Much like the outer membrane of Gram-negative bacteria, the MM contains integral proteins that we refer to as mycolate outer membrane proteins (MOMPs) (**Fig. 1A**) (4–8). In fast-growing mycobacteria such as *Mycobacterium smegmatis*, nutrient uptake across the MM is facilitated by Msp family proteins (9, 10). Of the known mycomembrane proteins, Msp complexes most closely resemble Gram-negative beta-barrel proteins. However, unlike traditional beta-barrels, individual Msp protein subunits contribute beta-strands to a larger beta-barrel (11). Msp family proteins are not conserved beyond fast-growing mycobacteria (12).

Another class of MOMPs found in slow-growing mycobacteria (*M. tuberculosis*, *M. avium*) are the proline-glutamate/proline-proline-glutamate (PE/PPE) proteins (13, 14). These proteins contain small alpha-helical N-terminal domains of approximately 100 (PE) or 180 (PPE) amino acids with diverse C-termini ranging in size from a few additional amino acids to large domains (>1000 residues). PE/PPE pairs are often functionally linked and encoded together in the same genetic locus. They typically require the type VII secretion system (Esx) for export, and in slow-growing mycobacteria they have been implicated in virulence and nutrient acquisition. Some PE/PPE proteins are thought to be associated with the surface of the mycomembrane (15, 16) or secreted into the extracellular milieu (17), while others have been proposed to form membrane channels across the mycomembrane for nutrient acquisition (7, 18–21).

However, the mechanism by which these proteins are integrated within the membrane and whether their insertion is coordinated with other aspects of envelope biogenesis has remained unclear. Although corynebacteria do not encode identifiable PE/PPE proteins, they are also thought to use pairs of small alpha-helical proteins that function as porins in the MM for nutrient transport (4, 22–27). These porins undergo a unique post-translational modification called *O*-mycoloylation in which proteins are modified with mycolic acid(s) by the periplasmic mycoloyltransferase Cmt1 (Cgp_0413) (28–31), and Cmt1 is required for their assembly into the MM (29, 31–34).

Besides the Sec translocon, members of the Mycobacteriales lack homologs of the known outer membrane protein assembly factors of Gram-negative bacteria, like the beta-barrel assembly machine (BAM complex) (35). Thus, the MOMP assembly pathway is likely to be distinct from that employed by Proteobacteria and other organisms with related diderm envelope architecture. To investigate these mechanisms, we have been using *Corynebacterium glutamicum* (*Cglu*) and its porin protein PorH (Cgp_3009) as a model system (32). PorH is encoded in an operon with another porin called PorA (Cgp_3008) (24). We show that PorA is required for the surface exposure of PorH and that *O*-mycoloylation of at least one of the porins plays a role in MOMP transport and assembly. Unlike for PE/PPE proteins, the type VII secretion system of *Cglu* is not required for their surface assembly. To investigate the spatiotemporal control of porin insertion into the MM, we probed the surface exposure of ALFA-tagged variants of PorH and another mycomembrane protein ProtX (Cgp_2875) (28, 29, 31) with fluorescently labeled anti-ALFA nanobodies. This analysis detected protein insertion at the cell poles that coincided with zones of active PG biogenesis. Moreover, fluorescence recovery after photobleaching (FRAP) of labeled MOMPs and extraction of the mycoloyl arabinogalactan-peptidoglycan (mAGP) sacculus revealed that PorH and ProtX are immobile in the MM and likely attached to the cell wall. Genetic and biochemical assays reported here and in the literature suggest that PorH and PorA form a hetero-oligomeric pore (22). Consistent with this, AlphaFold2 (36, 37) predicts a high confidence structure of a hetero-oligomeric pore formed by PorA and PorH. Overall, our results indicate that like Gram-negative bacteria, *Cglu* and potentially other members of the Mycobacteriales order coordinate outer membrane protein assembly with PG biogenesis, have immobile outer membrane proteins, and have a physical connection between the outer membrane and the cell wall through outer membrane proteins.

## RESULTS

### PorH requires PorA and *O*-mycoloylation but not the type VII secretion system for its surface assembly

PorH is a small protein (57 amino acids/6.1kDa) predicted to adopt an alpha-helical structure. It is encoded upstream of a similarly small (45 amino acids/4.7kDa) protein called PorA (24) that is also predicted to be alpha-helical in structure. Inactivation of either PorH or PorA in *Cglu* results in increased kanamycin resistance, suggesting that these proteins promote cell permeability, possibly by forming a pore structure in the MM (**Fig. 1B**) (23, 32). Accordingly, a 6x-His tag fused to the C-terminus of PorH (PorH-His) is surface accessible (32, 38), and *in vitro* experiments with non-native lipid membranes detected channel forming activity for mixtures of purified PorA and PorH (22). Thus, PorA and PorH as well as other related porin protein pairs of corynebacteria (4) are thought to function as two-component channel forming complexes in the mycomembrane.

Given the potential for PorH to form a complex with PorA in the MM, we tested whether PorA is required for PorH surface assembly. Also, because *Cglu* encodes homologs to some *M. tuberculosis* type VII secretion system proteins that may comprise one or more functional systems (39), we investigated whether these porins might be transported through the type VII secretion system like the PE/PPE proteins in slow-growing mycobacteria (13). Proper assembly of PorH in the MM was assessed by monitoring surface accessibility of the 6x-His tag of a PorH-His fusion to a fluorescently labeled anti-His antibody. Upon deletion of *porA*, the normally surface-exposed tag of PorH-His was not detected by the antibody as assessed by flow cytometry (**Fig. 1C**). Surface accessibility of the tag was restored upon production of PorA from an ectopic locus in a Δ*porA* strain (**Fig. 1C**). Deletion of genes encoding homologs of type VII secretion components *eccB3* (*cgp_0661*), *eccC5* (*cgp_2498*), or *eccD2* (*cgp_0830*) did not reduce surface exposure of PorH-His (**Fig. S1A**), indicating that PorH is unlikely to be a type VII secretion substrate.

Notably in the absence of PorA, anti-His immunoblots detected PorH-His accumulation in cells with the protein appearing in forms that migrate more slowly on an SDS-PAGE gel (**Fig. 1D**). The cause of the altered mobility is not known. However, the buildup of PorH-His that is not surface exposed upon *porA* deletion is reminiscent of unfolded, “off-pathway” outer membrane proteins that accumulate in the *Escherichia coli* periplasm when there are defects in the outer membrane protein assembly pathway. Such defects occur when there are mutations in the coding sequence of the outer membrane protein itself that inhibit transport and/or assembly or upon inactivation of quality control/transport factors involved in outer membrane protein transport, such as chaperones (40–42). The buildup of PorH-His that we observe in the absence of PorA may represent a similar accumulation of protein that has “fallen off” of the assembly pathway.

Another factor known to be required for PorH surface assembly is the protein mycoloyltransferase Cmt1 (Cgp_0413) (**Fig. S1B**) (28, 30–32). Cmt1 has many substrates (29, 33, 34). Thus, it is unclear if its role in PorH surface assembly involves *O*-mycoloylation of the porin proteins themselves, the *O*-mycoloylation of a critical assembly factor, or both. We therefore investigated the effect of amino acid substitution at the *O*-mycoloylation sites of PorA and PorH on the surface assembly of PorH-His. Substitution of the *O*-mycoloylated serine residues in either PorH (S56A) or PorA (S15A) did not affect the surface exposure of the His tag or the function of the pore as assessed by kanamycin sensitivity (**Fig. 1E and Fig. S2**). These results indicate that *O*-mycoloylation of either PorH or PorA is sufficient for transport and function of the porin.

Although substitutions affecting *O*-mycoloylation of a single porin did not affect PorH assembly, surface detection of PorH-His was abolished when the *O*-mycoloylation of both proteins was blocked by the serine substitution (**Fig. 1E**). The cells producing PorH^S56A^-His and PorA^S15A^ also displayed increased kanamycin resistance, indicating the failure to form a functional complex (**Fig. S2**). Notably, unlike cells lacking Cmt1 that fail to accumulate PorH-His whether or not PorA is produced (**Fig. 1D and 1F**) (32), PorH^S56A^-His accumulated in what appear to be “off pathway” forms in cells co-producing the PorA^S15A^ variant (**Fig. 1F**). From these results, we conclude that PorH assembly into the MM requires PorA and the *O*-mycoloylation of either itself or PorA, providing support for the idea that the two proteins assemble and function as a complex. Furthermore, because PorH-His mobility is affected by its mycoloylation status or the mycoloylation status of PorA, we infer that *O*-mycoloylation is required for the normal PorH maturation process.

### PorH is inserted into the mycomembrane at zones of active cell envelope growth

We next investigated the subcellular location of PorH assembly at the cell surface. For these studies, we used an ALFA-tagged version of PorH produced from an expression construct under the control of an IPTG-inducible *tac* promoter (P*_tac_*::*porH-ALFA*) in a strained deleted for the native *porH* gene. The functionality of the PorH-ALFA fusion was confirmed by its ability to restore wild-type kanamycin sensitivity in the *porH* deletion strain, indicating that the ALFA tag does not interfere with PorH function (**Fig. S3**). To visualize the assembly of PorH at the cell surface, exponentially growing cells were immobilized on an agarose pad supplemented with IPTG (500μM) and Atto488-conjugated anti-ALFA nanobody (Atto488-NB, 12.5nM) and the appearance of fluorescent signal was monitored over time. We found that adding a dilute concentration of Atto488-NB to the agarose pads yielded more uniform labeling than incubating the cells with the nanobody in solution and washing away unbound molecules before imaging on agarose pads (**Fig. S4A**). PorH signal was enriched at the old cell poles and at division sites following the rapid process of septum splitting called V-snapping (new poles) (**Fig. 2A-B, Movie S1**) (43, 44). This labeling pattern remained consistent over a broad range of IPTG concentrations (50 to 500μM) as did the correlation of PorH-ALFA incorporation with the rate of cell elongation (**Fig. S4C and S5**). Moreover, the rate at which labeled PorH-ALFA transitioned from the pole to the sidewall following a chase without Atto488-NB was also unaffected by the level of induction (**Fig. S5A-D**). Importantly, expression of untagged PorH did not result in any detectable fluorescence, confirming the specificity of the observed labeling pattern (**Fig. S4D**).

**Figure 2:**
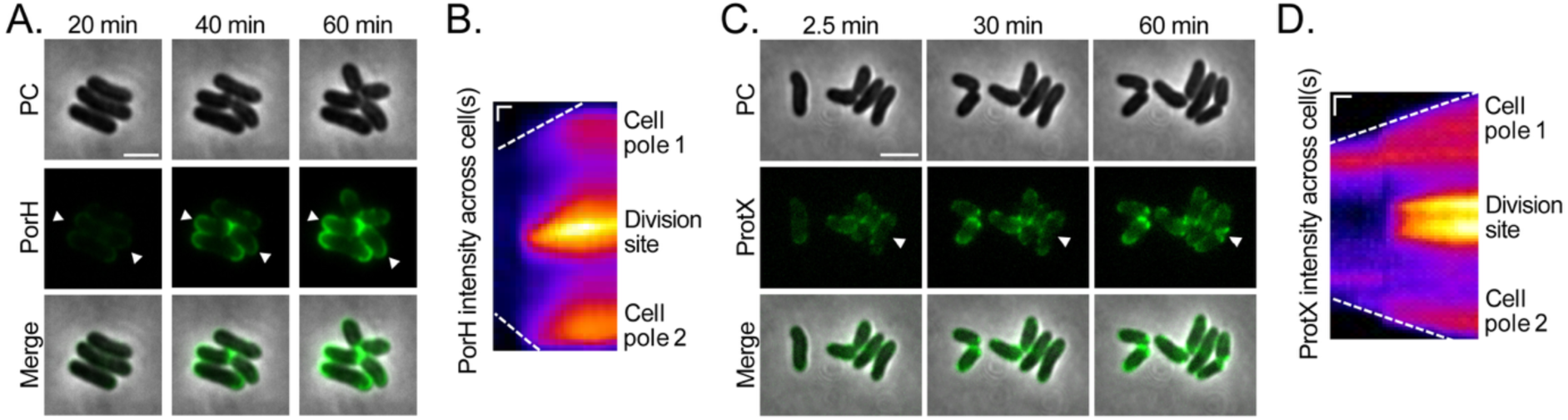
PorH and ProtX surface assembly occurs at old and new cell poles. A representative image of cells expressing **(A)** PorH-ALFA (strain H1616) and **(C)** ProtX-ALFA (strain DB42) at indicated timepoints after they were spotted on agarose pads supplemented with 500 µM IPTG and 12.5 nM Atto488-NB. See *Methods* for details. Scale bar = 3 µm. **(B and D)** Kymograph of the cell highlighted with arrowheads in **(A and C)** throughout the entire time-lapse, showing the **(B)** PorH-ALFA and **(D)** ProtX-ALFA incorporation over time (horizontal axis). Vertical scale bar = 0.5 µm, Horizontal scale bar = 10min. Data is representative for biological triplicates and in total n = 113 cells for PorH-ALFA and n = 1270 cells for ProtX-ALFA were imaged.

In addition to PorH, we were also able to detect surface labeling of another known *Cglu* MOMP called ProtX (28, 29, 31). We applied the same labeling strategy as for PorH by expressing a C-terminal ALFA-tagged version of ProtX and detecting its surface exposure as described above. Similar to PorH, we observed preferential incorporation of ProtX at the old and new cell poles, the latter formed after V-snapping (**Fig. 2C-D, Fig. S4B, Movie S2**). As with PorH, staining was entirely dependent on the expression of the ALFA-tagged construct and remained consistent across a wide range of inducer concentrations (**Fig. S4E-F**).

To correlate MOMP membrane incorporation with overall cell envelope biogenesis, we followed PorH-ALFA surface exposure alongside labeling with 7-hydroxycoumarin-3-carboxylic acid–D-alanine (HADA) as a marker for PG synthesis (45) and the localization of the polar scaffolding protein DivIVA fused to mScarlet (46). Although HADA is incorporated by both PG synthases (D,D-transpeptidases) and remodeling enzymes (L,D-transpeptidases) (47), the labeling pattern correlates well with nascent PG synthesis as monitored using the D-Ala-D-Ala probe EDA-DA that is incorporated into the PG layer as modified precursors that can be labeled with click-chemistry compatible dye reagents (**Fig. S6**) (48). Because the appearance of PorH-ALFA fluorescence at the poles following IPTG addition involves many steps (transcription, translation, export, etc), there is a time-delay for PorH-ALFA labeling relative to the rapid incorporation of HADA label. Therefore, a two-color, pulse-chase labeling approach was used to follow PorH-ALFA assembly (**Fig. 3A**). In a microfluidic device (CellASIC), PorH-ALFA expression was induced, and cells were pre-labeled with Atto488-NB for 75 minutes (approximately 1.5 generations). The medium was then switched to remove free Atto488-NB and flow in Alexa647-conjugated anti-ALFA nanobody (Alexa647-NB) was added along with HADA. With this approach, newly incorporated PorH-ALFA labeled with Alexa647-NB can be directly compared with PG biogenesis detected by HADA labeling (**Fig. 3B**). As observed previously, newly incorporated Alexa647-NB labeled PorH-ALFA localized to the cell poles (**Fig. 3B**). Moreover, segmentation of 400 cells across three biological replicates in combination with analysis of their fluorescence intensity profiles over time further confirmed this incorporation pattern and demonstrated that nanobody binding is compatible with live-cell imaging in the microfluidic setup (**Fig. 3C**).

**Figure 3:**
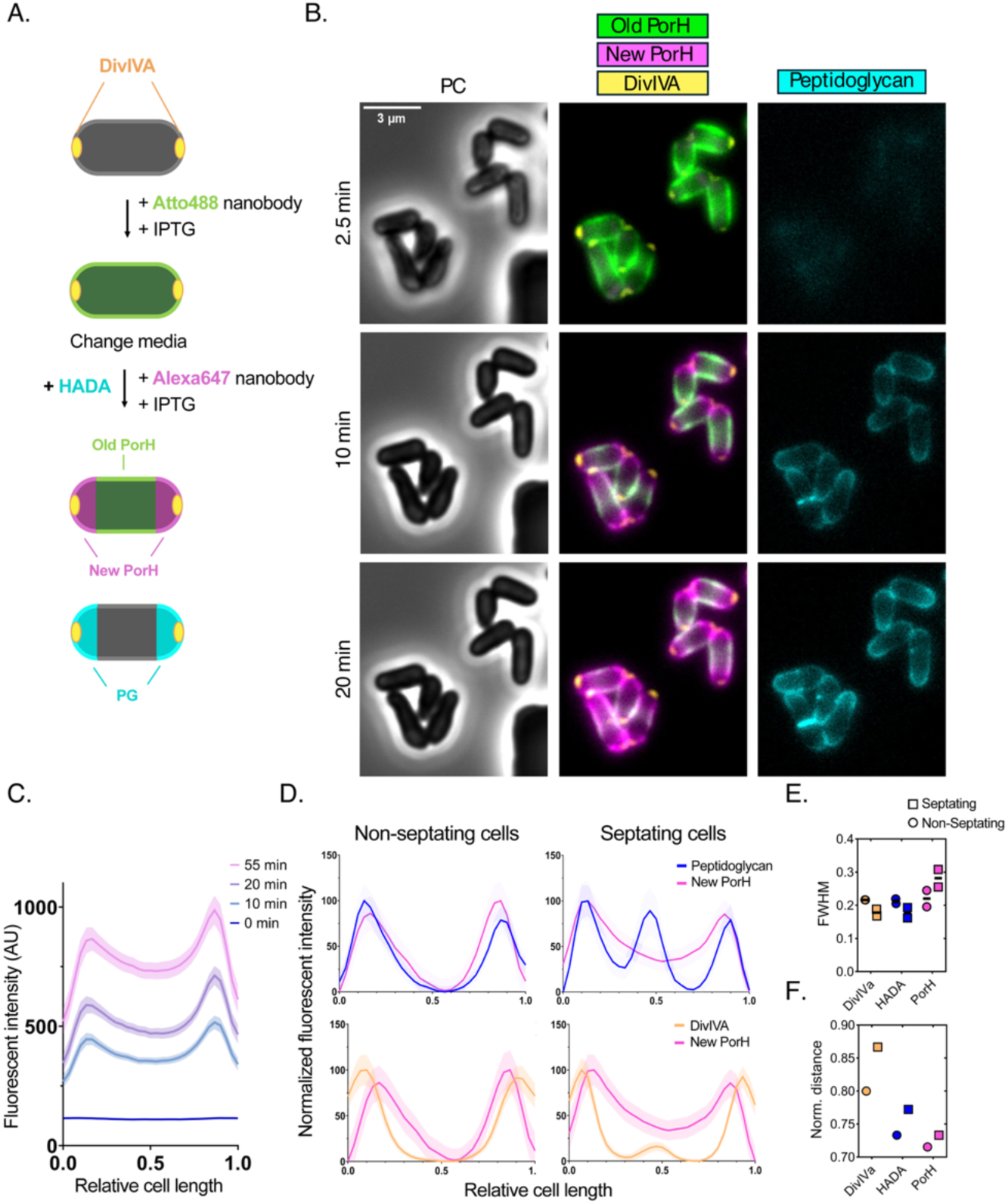
PorH is incorporated at zones of active cell wall synthesis. **(A)** Schematic representation of pulse-chase experiment using microfluidics. Note that cells (strain DB16) were chased with Alexa647-NB and HADA simultaneously after the initial labeling with Atto488-NB labeling. The schematics for Alexa647-NB and HADA labeling are shown separately for clarity. See main text for further details. **(B)** A representative image from biological triplicates of the microfluidic pulse chase experiment at indicated time points. Scale bar = 3 µm. **(C)** Quantification of Alexa647-NB labeled PorH fluorescence intensity as a function of normalized cell length at indicated time points following IPTG induction. Cells were oriented based on highest HADA intensity (= Pole 0). A representative graph from one biological replicate of n = 238 cells is shown, shaded area indicated ±95% confidence interval. **(D)** Comparison of normalized fluorescence intensity profiles across cell length for PG (blue), PorH (magenta), and DivIVa (orange) in non-septating and septating cells. Cells were oriented based on HADA intensity. n = 494, plotted from 3 replicates, shaded area indicates ±95% confidence interval. **(E and F)** Full width at half max (FWHM) values **(E)** and the distance between polar peak centroids **(F)** for the fluorescent intensity profiles in **(D)**.

Next, we compared the fluorescence intensity profiles of PorH-ALFA to those of HADA and DivIVA after a 10-minute chase with Alexa647-NB. As previously shown, DivIVA displayed the most sharply defined polar localization, while both HADA and PorH-ALFA showed broader distributions extending from the polar cap into the cylindrical cell body (**Fig. 3D**) (46). We further classified cells as either septating or non-septating, based on the presence or absence of HADA signal at mid-cell. In non-septating cells, PorH-ALFA and HADA signals were strongly correlated (Pearson’s correlation coefficient, r = 0.9025) (**Fig. S7**). In septating cells, however, the correlation was lower (r = 0.6011) due to the absence of PorH-ALFA incorporation at the septum prior to V-snapping as well as the broader polar distribution of PorH-ALFA compared to HADA. One limitation of this assay is that the anti-ALFA nanobody can only detect surface exposed epitope tags and thus likely cannot reach the inward growing septum. However, despite the lack of a defined septal band (as observed for HADA), septating cells showed overall higher fluorescence intensity of PorH-ALFA at mid-cell compared to non-septating cells (**Fig. 3D**). Taken together, our imaging data suggest that MOMPs, such as PorH and ProtX, are incorporated at sites of active cell envelope biogenesis. However, the zone of MOMP incorporation appears to extend slightly into the sidewall (**Fig. 3E**), and the fluorescence peaks of MOMP signals are not quite as separated as those of PG (**Fig. 3F**), suggesting that MOMP insertion can occur as the polar envelope material matures into a sidewall structure. Moreover, as indicated by the clear separation of old and newly labeled PorH, MOMPs appear to remain stably embedded once incorporated into the MM and are slowly displaced toward the sidewall by the insertion of new cell envelope material at the pole (**Fig. 3B**).

### PorH is non-diffusive in the mycomembrane

In Gram-negative bacteria, outer membrane proteins show low diffusivity due to the strong lateral interactions between lipopolysaccharide molecules in the outer leaflet of the outer membrane (49–53). We were therefore curious to know whether PorH is diffusible in the MM, especially considering that unlike the outer membrane of Gram-negative bacteria, glycolipids in the MM of *Cglu* are diffusive (54). To investigate the diffusion dynamics of PorH within the MM, we employed fluorescence recovery after photobleaching (FRAP) of cells producing PorH-ALFA labeled with Atto488-NB. As controls, we measured FRAP of MM lipids labeled with the fluorescent trehalose analog 6TMR-tre (54) and PG labeled with HADA (45). In agreement with previous studies, 6TMR-tre fluorescence was diffusive, exhibiting fluorescence recovery on the scale of seconds (**Fig. 4A-B**) (54). Labeled PG showed no recovery as expected from a covalently crosslinked matrix (**Fig. 4A-B**). Labeled PorH-ALFA was bleached within the cell cylinder and on the cell poles. No fluorescence recovery was observed in the timescale of the experiment at either location (1min total) (**Fig. 4A-B, Movie S3**), demonstrating that PorH is non-diffusive within the diffusive MM over tens of seconds.

**Figure 4:**
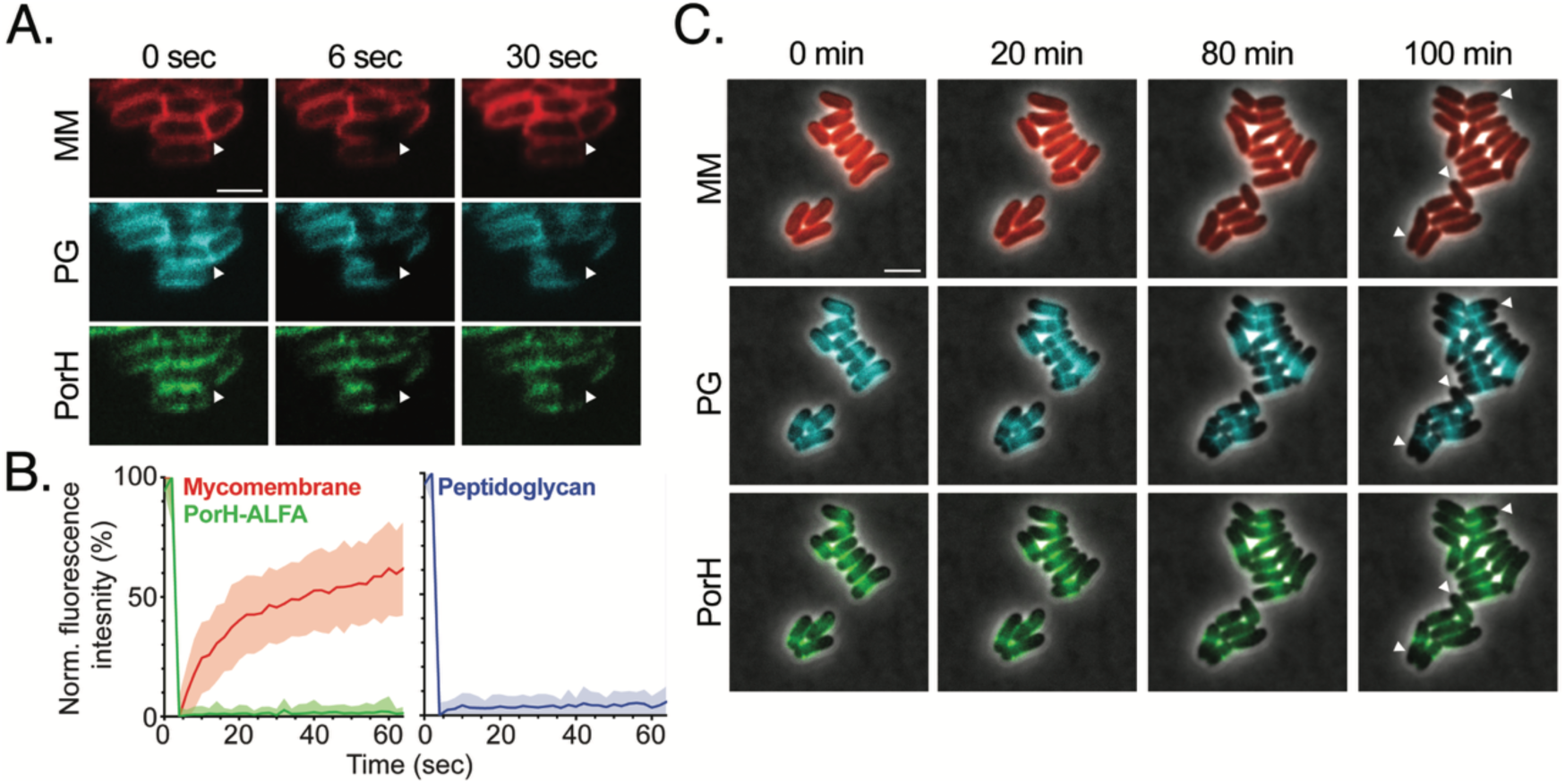
PorH is non-diffusive in the mycomembrane. **(A)** A representative image and corresponding **(B)** FRAP curves of n = 35 cells from biological duplicates (strain H1616) labeled with 30 µM 6-TMR-tre (MM), 100 µM HADA (PG) and 12.5 nM Atto488-NB (PorH-ALFA) (see *Methods* for details). Arrowheads indicate bleached regions. Scale bar = 2 µm. **(C)** Fluorescence dilution assay for PorH-ALFA. Cells (strain H1616) were prelabeled with same dyes as in **(A)** and imaged over 4h after dye removal with a 20min acquisition frame rate. Arrowheads indicate regions of new polar insertion (unlabeled material formed after dye was removed). Scale bar = 3 µm.

We additionally assessed the diffusion of PorH-ALFA over longer timescales (tens of minutes to hours) using a fluorescence dilution assay. To this end, cells were fully pre-labeled with 6-TMR-tre, HADA, and Atto488-NB before being placed on an agarose pad in the absence of label for over 4 hours. The signal for the 6TMR-tre in the MM decreased uniformly in intensity as cells grew (**Fig. 4C, Fig. S8, Movie S4**). However, the HADA label in PG and the Atto488-NB label on PorH-ALFA showed no detectable redistribution during the extended growth period, retaining high Pearson’s correlation coefficients (r > 0.81), indicating that these components remain stably localized in contrast to the mycomembrane (**Fig. 4C, Fig. S8, Movie S5-6**). We also performed the FRAP and fluorescence dilution assays for ALFA-tagged ProtX. As was seen for PorH, no fluorescence recovery was observed after photobleaching, and the fluorescent signal remained static during outgrowth (**Fig. S8**). Overall, our results indicate that PorH and ProtX remain fixed in place after insertion into the MM despite the membrane itself being diffusive.

### PorH and ProtX are linked to the cell wall

One potential explanation for the non-diffusive behavior of PorH and ProtX in the MM is that they are attached to the cell wall (AG and/or PG). To test this possibility, we purified the mAGP (mycoloyl-arabinogalactan-peptidoglycan) fraction of the cell envelope from cells expressing untagged or ALFA-tagged versions of the MOMPs. The mAGP fraction contains the PG sacculus with linked AG and any mycolic acid that is covalently attached to the AG (55). Without further treatment, proteins in the mAGP were unable to enter an SDS-PAGE gel, presumably due to their covalent or noncovalent association with the cell wall (**Fig. 5A and 5C**). However, lysozyme treatment liberated proteins from the mAGP fraction, allowing for the detection of Coomassie stainable bands in the gel (**Fig. 5A and 5C**). A similar lysozyme treatment of mAGP isolated from *M. smegmatis* failed to liberate proteins, indicating that the mAGP from *Cglu* is more permissive to fractionation with lysozyme than *M. smegmatis* mAGP (**Fig. S9**), presumably due to the shorter mycolic acid chains covalently attached to the AG in corynebacteria relative to mycobacteria (56). Immunoblot analysis of the mAGP fractions from *Cglu* detected PorH-ALFA or ProtX-ALFA only in the tagged samples digested with lysozyme (**Fig. 5B and 5D**). The mAGP extraction protocol includes both sonication and boiling in a high concentration of detergent (55). Similar extraction protocols have been used to remove soluble proteins from cell envelope preparations of *M. tuberculosis* (57), to isolate the mAGP for downstream mass spectrometry analysis (55, 58, 59), and to analyze proteins associated with the mycobacterial cell wall (60). Thus, the material released from the mAGP fractions of *Cglu* by lysozyme are unlikely to represent noncovalently associated proteins trapped within the wall matrix. Further support for the covalent linkage of PorH to the cell wall comes from the observation that it is released from the mAGP fraction upon trifluoroacetic acid (TFA) treatment (**Fig. S10B**) (61–63). Notably, TFA treatment does not lead to the bulk release of protein from the mAGP like lysozyme treatment (**Fig. S10A**), but such a release remains possible following lysozyme digestion of the TFA-treated sacculi (**Fig. S10C**). Based on these fractionation experiments and the observed immobility of the porins in the envelope of live cells, we conclude that PorH and ProtX are likely to be covalently linked to the cell wall.

**Figure 5:**
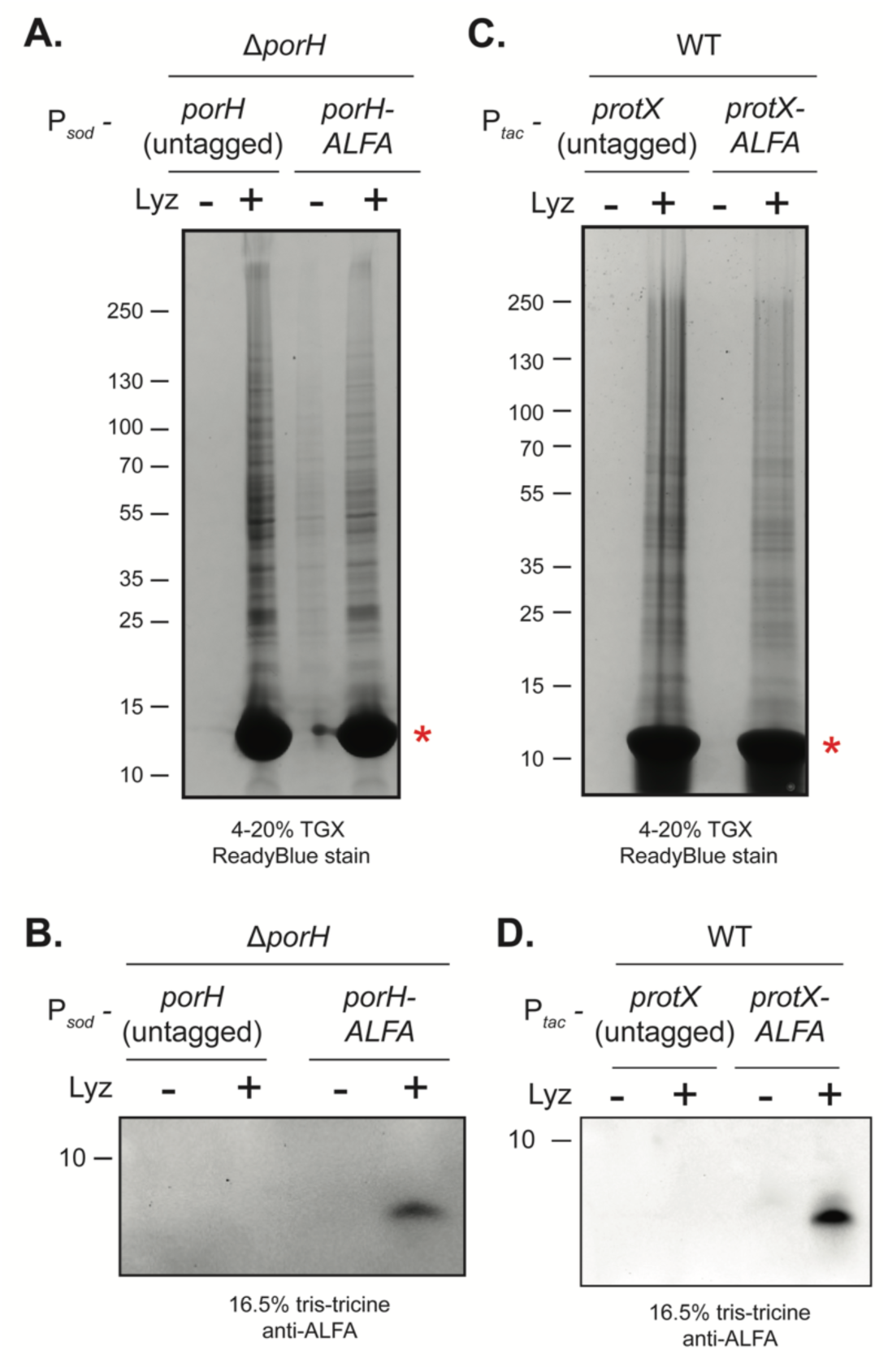
PorH and ProtX are linked to the cell wall. The mycoloyl-arabinan-peptidoglycan (mAGP) cell wall fraction was isolated from cells expressing either **(A-B)** untagged PorH or PorH-ALFA from a constitutive P*_sod_* promoter in a Δ*porH* background or **(C-D)** untagged ProtX or ProtX-ALFA from a P*_tac_* inducible promoter in a wild-type (WT) background expressed with 1mM IPTG. The resulting mAGP fraction was left undigested or digested with 5mg/mL lysozyme and analyzed by SDS-PAGE followed by ReadyBlue protein staining **(A and C)** and immunoblot analysis **(B and D)**. Red asterisk indicates the band corresponding to the added lysozyme. All experiments were performed in biological triplicate and a representative replicate is displayed.

To determine if the association of the porins with the mAGP is dependent on a properly formed AG layer (64–67), we performed fractionation experiments in cells lacking arabinosyltransferases involved in AG biogenesis (AftB, AftC, AftD) or cells treated with sub-MIC levels of the AG synthesis inhibitors TCA1 or EMB. Deletion of arabinosyltransferases responsible for AG branching (AftC and AftD) or for adding the terminal sugars (AftB) did not impact levels of PorH-ALFA detected in whole cells (**Fig. S11C**) or linked to the mAGP (**Fig. S11A-B**). However, both TCA1 and EMB, which inhibit early stages of AG biosynthesis (2, 68–70), led to a strong reduction in PorH-ALFA detected in the mAGP fraction (**Fig. S11A-B**) without affecting the amount detected in whole cells (**Fig. S11C**). Thus, AG synthesis is required for the stable association of PorH with the mAGP fraction, suggesting that it may be linked to the AG layer.

### Model structure of the PorA-PorH pore complex

How PorH might form a pore in the MM in complex with PorA has remained unclear. Given the small size of each protein, it is likely they are assembling into an oligomeric complex to generate a pore structure. We leveraged advancements in protein prediction algorithms to predict the structure of the PorH/PorA porin complex from *Cglu* with AlphaFold2 (36, 37). Predictions with an increasing number of subunits of each protein were made (**Fig. S12**). The prediction that formed a pore structure with high confidence was the a hetero-pentameric structure containing five copies each of the PorH monomer and PorA monomer (pLDDT = 94.5, pTM = 0.889, ipTM = 0.883) (**Fig. 6 and S12**). In this predicted structure, monomers of PorH and PorA reside in antiparallel orientation and are twisted to form a tornado-like pore (**Fig. 6A-B**). Several features of this structure provide confidence in its potential biological relevance. First, the *O*-mycoloylation sites, PorH^S56^ and PorA^S15^ (28, 31), form a ring around the structure that would create a belt of lipidated residues to embed the complex within the MM (**Fig. 6C**), much like the so-called “aromatic girdle” of Gram-negative β-barrel outer membrane proteins (71–73). Second, the outside of the structure which would face into the MM lipid bilayer, is hydrophobic whereas the inside of the pore is hydrophilic (**Fig. 6A**). Third, the constricted pore size of the predicted structure is roughly 30Å, which is similar to the pore diameter of β-barrels in *E. coli* such as OmpF (74) and OmpX (**Fig. 6A**) (75). Finally, the C-terminal residues of PorH are located at the outer “rim” of the pore, explaining surface accessibility of this terminus (**Fig. 6C**) (32, 38).

**Figure 6:**
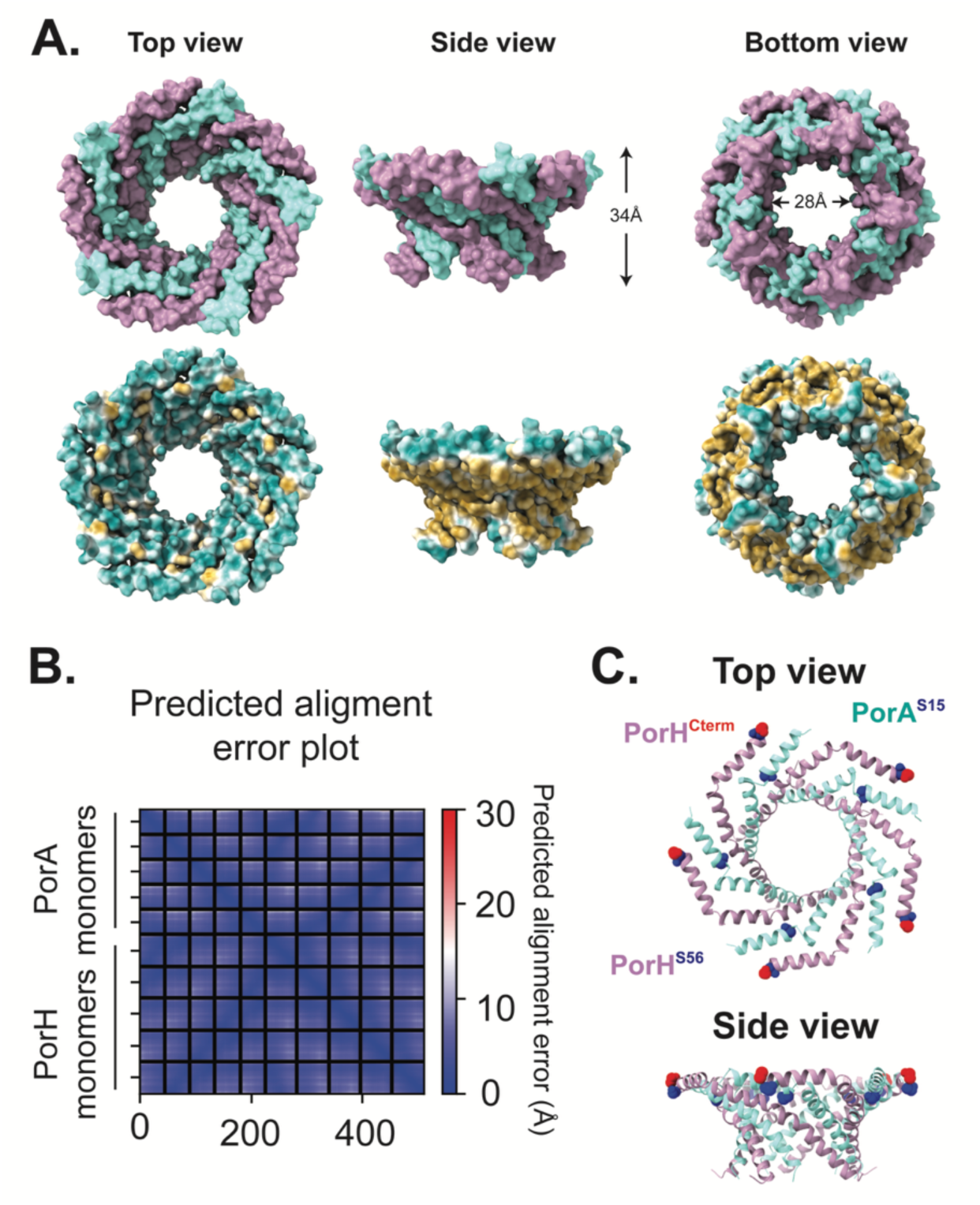
Model of the PorAH pore. The predicted structure was generated with AlphaFold multimer (36, 37). All structure images and measurements were generated or visualized using UCSF Chimera (110). **(A)** AlphaFold multimer structural prediction of PorAH hetero-pentamer. The top row of images is colored with by protein (PorA in cyan and PorH in purple). The bottom row of images is colored by hydrophobicity with yellow indicating more hydrophobic residues and cyan indicating more hydrophilic residues. Measurement of the height of the predicted complex was made between residues V13 and G45 on PorA. Measurement of the predicted pore size was made between residue E15 and N19 on PorH. **(B)** Predicted alignment error (PAE) plot of the AlphaFold2 predicted structure showing the positional error for each residue with blue representing less predicted error and red indicating more predicted error. **(C)** Top and side views of the predicted structure in ribbon view. *O*-mycoloylation residues on PorH and PorA, S56A and S15A, respectively, are indicated by blue spheres. The C-terminus of PorH (S57) is indicated by red spheres.

## DISCUSSION

Despite their importance for nutrient uptake, virulence, and drug sensitivity, the pathways for protein transport and integration within the MM of corynebacteria and mycobacteria remain poorly characterized. To better understand this process and how it is coordinated with other aspects of bacterial envelope assembly, we have investigated surface assembly of the MOMPs PorH and ProtX in *Cglu* as a model system. For PorH, we found that surface assembly requires its partner porin protein PorA and the *O*-mycoloylation of at least one member of the pair by the mycoloyltransferase Cmt1 (**Fig. 1C, 1E**). Furthermore, we showed that the assembly of both PorH and ProtX within the MM occurs at the polar growth zones coincident with PG biogenesis (**Fig. 2**). Finally, we demonstrate that the diffusion of these proteins is limited once they are assembled on the surface and we propose that this is due to their linkage to the cell wall (**Fig. 4**). These findings reveal that although corynebacteria and Gram-negative bacteria like *E. coli* have outer membranes that differ fundamentally in structure and composition, their surface protein assembly processes share similar properties.

### The porin assembly pathway

In Gram-negative bacteria, outer membrane protein assembly is divided into distinct stages: secretion across the plasma membrane by the Sec translocon, transport through the periplasm by chaperones, and folding within the outer membrane by the BAM complex. Notably, the porins PorA, PorH, and ProtX lack identifiable signal sequences for Sec transport (31). Thus, in addition to lacking a type VII secretion system requirement (**Fig. S1**), the mechanism by which these porins cross the plasma membrane remains unclear.

Once they transit the plasma membrane, the porins likely interact with the mycoloyltransferase Cmt1 and for PorH, its binding partner PorA. We and others have shown that PorH levels are drastically reduced on both the cell surface and in whole cell lysates when Cmt1 is inactivated (**Fig. 1D, Fig. S1B**) (29, 31, 32). PorH is similarly absent from the cell surface when its binding partner PorA is inactivated or the mycoloylation sites of both PorA and PorH are inactivated (**Fig. 1C**, **Fig. 1E**). By contrast, there is an accumulation of intracellular PorH in these genetic backgrounds (**Fig. 1D**, **Fig. 1F**). Thus, the intracellular buildup of PorH is dependent on Cmt1, but not on *O*-mycoloylation. This finding is reminiscent of the accumulation of outer membrane proteins in the periplasm of *E. coli* upon inactivation of periplasmic chaperones like SurA (41, 76, 77). Therefore, although it remains possible that Cmt1 indirectly controls proteolysis of PorH through *O*-mycoloylation of a periplasmic protease, we favor a model in which Cmt1 not only lipidates PorH but also functions as a chaperone for the porin during its transport across the periplasm to the cell surface. Accordingly, Cmt1 has been detected in both the soluble and membrane fractions of *Cglu* and thus could serve as a shuttle between compartments (78–80). Additionally, a recent crystal structure of Cmt1 with an analogue of the mycomembrane lipid trehalose monomycolate has been solved, shedding light on how Cmt1 could accommodate the *O*-mycoloylation modification of PorH (34).

In Gram-negative bacteria, insertion of beta-barrel proteins into the outer membrane by the BAM complex is coordinated with PG synthesis (81, 82). Similarly, we observe that the surface assembly of PorH and ProtX occurs at sites of polar PG biogenesis (**Fig. 4**). What remains unclear is whether secretion of the porins across the plasma membrane also occurs at the poles or if secretion occurs at dispersed locations around the cell periphery followed by targeting to the poles for surface assembly. It also remains unknown whether a specific machinery is required for MOMP insertion or whether surface assembly is spontaneous but somehow restricted to sites of new envelope growth.

In addition to the polar sites of assembly, we also observe MOMP surface assembly at division sites (**Fig. 3**). However, the process appears to be limited to the later stages of division. Formation of the septum in *Cglu* proceeds through temporally distinct steps (54). First, the new PG and AG layers are formed, followed by the influx of mycolic acids into the septum. This stage of MM formation is then closely followed by a rapid cell separation process called V-snapping. Our data suggest that MOMP incorporation begins immediately following V-snapping, the first point at which the septal region becomes exposed to the extracellular environment. Because our labeling approach relies on surface accessibility, it remains uncertain whether the timing of MOMP assembly at the division site reflects a true delay in MOMP insertion at the septum or a limitation of the detection method. However, given that both PorH and ProtX are *O*-mycoloylated, we favor a model in which they are inserted during or immediately after MM assembly, which coincides with detection of surface exposure using nanobodies. Notably, regardless of the timing, only fully matured regions of the cell surface appear to be exposed to the environment upon V-snapping similar to Gram-positive organisms like *Staphylococcus aureus* that also divide via a rapid cell separation event (83).

### Porins are linked to the cell wall

In Gram-negative bacteria, outer membrane proteins have been shown to assemble into crystalline-like protein islands that display a confined lateral mobility (49, 51, 84, 85). This lack of diffusivity is due to lipopolysaccharide molecules that make up the outer leaflet of the outer membrane. These lipids have been shown to be largely non-diffusive due to lateral packing interactions and interactions with outer membrane proteins (50, 51). In contrast to the Gram-negative outer membrane, lipids within the MM of *Cglu* have been reported to exhibit diffusive behavior (54). It was therefore surprising to observe that PorH and ProtX do not display diffusive behavior in the MM but instead appear to be immobile like the outer membrane proteins of Gram-negative bacteria (**Fig. 4**). Additionally, like the Gram-negative beta-barrel proteins, our results from pulse-chase and fluorescence dilution experiments demonstrate that there is little to no turnover of MOMPs once they are inserted into the MM (**Fig. 3**).

How is protein localization confined within a fluid membrane? Our detection of PorH and ProtX in the mAGP fraction indicate that the spatial constraints on PorH and ProtX diffusion are likely to arise from their linkage to the cell wall (**Fig. 5**). Lysozyme digestion or trifluoroacetic acid hydrolysis of the mAGP (**Fig. 5, Fig. S10**) liberates porins from the mAGP, and their association with the cell wall is dependent on proper synthesis of the AG layer (**Fig. S11**). This observation coupled with the proximity of the AG layer to the MM (64–66) suggests that the porins are linked to the AG, but this possibility requires further investigation.

In Gram-negative bacteria, covalent linkage of outer membrane-anchored proteins like Lpp to the PG is critical for envelope structure and integrity (86–89). In corynebacteria and mycobacteria, a subset of the mycolic acids that form the MM are covalently linked to the AG layer (90), providing a similar outer membrane-cell wall connection. Given that such connections are already in place, it remains unclear why MOMP-cell wall connections might also be necessary. One possibility is that they provide additional MM-cell wall connections to help stabilize the envelope. Accordingly, *Cglu* mutants inactivated for AftB, which adds the terminal sugar on AG needed for the covalent attachment of mycolates (79, 91), shed MM fragments but do not completely lack a MM. Thus, the MOMP-cell wall connections may provide a second set of MM-AG linkages that together with the mycolate-AG bonds support the formation of a stable MM.

### Possible pore structure?

How the small alpha-helical porin proteins like PorA and PorH form a pore in the MM has remained unclear for some time. Although biochemical and structural analysis is required, the AlphaFold2 prediction of the PorH-PorA hetero-pentamer provides an attractive model for pore formation by such small proteins. One feature of the predicted pore structure that is difficult to rationalize is its depth (∼34Å). Cryo-sections of *Cglu* cells show variability in the width of the MM, ranging from 50Å to 80Å (64–66, 92). The MM, then, may have sections that vary in width with thinner areas accommodating the PorAH pore.

Alternatively, the PorAH pore may exist in complex with other MOMPs or in a larger oligomeric complex, such as a dimer of hetero-pentamers, to span the MM. Further work will be needed to distinguish between these two possibilities.

### Potential parallels with mycobacterial MOMPs?

There are no known homologs of corynebacterial porin proteins encoded by mycobacteria. Instead, fast-growing mycobacteria utilize the Msp family of beta-barrel proteins to mediate nutrient uptake across the MM (93). Slow-growing mycobacteria, on the other hand, encode numerous PE/PPE proteins, some of which have been implicated in MM permeability, the transport of small molecules, and nutrient uptake (7, 14, 18, 94–97). Whether the PE/PPE proteins also form pores in the mycomembrane remains to be determined. However, this possibility seems likely given the genetic results and the properties they share with the corynebacterial porins such as their small, α-helical nature, cell surface accessibility, and propensity to be encoded within operons as pairs that are functionally interdependent (9, 13, 98).

Because of the diversity of MOMPs found among the Mycobacteriales, it is unclear whether proteinaceous links between the MM and cell wall will prove to be a general feature of these organisms or if this property is limited to corynebacterial porins. However, both lipoproteins and beta-barrel proteins have been implicated in providing connections between the outer membrane and cell wall in different diderm, Gram-negative bacteria (86–89). Thus, it is not unreasonable to expect that the MM-cell wall linkage observed here for the corynebacterial porins might be employed by mycobacteria and potentially more broadly among members of the Mycobacteriales.

## MATERIALS & METHODS

### Bacterial strains & growth conditions

All strains used in this study are listed in **Table S1**. All experiments were performed using *C. glutamicum* MB001, a prophage-free strain derived from ATCC13032 (99) and *M. smegmatis* mc^2^155. *M. smegmatis* strains were grown at 37°C in Middlebrook 7H9 media (BD) supplemented with 10% ADC (albumin, dextrose, catalase) and 0.05% Tween 80. *C. glutamicum* strains were grown in Brain Heart Infusion (BHI) medium (BD) or brain heart infusion medium supplemented with 9.1% sorbitol (BHIS) at 30°C with aeration, as indicated, unless harboring the pEWL89 cre recombinase plasmid or pEWL103 recombineering vector. Strains carrying these temperature-sensitive plasmids were propagated at 25°C. Ectopic expression constructs were induced using 1mM theophylline (for integrated riboE1 constructs) (100, 101) or 50-1000µM IPTG (for P*_tac_* replicating plasmids derived from pTGR5, 50-500µM for microscopy and 1000µM for mAGP detection) (102). *E. coli* DH5α (λ_pir_) or NEB10-beta (NEB) cloning strains were grown at 37°C in LB (1% tryptone, 0.5% yeast extract, 0.5% NaCl) with aeration unless harboring pCRD206 allelic-exchange derivatives, in which cells were cultured at 30°C. *C. glutamicum* strains were grown with the following antibiotics when appropriate: 15μg/mL kanamycin, 3.5μg/mL chloramphenicol, 12.5μg/mL apramycin, and 150μg/mL zeocin. *E. coli* strains were grown with the following antibiotics for plasmid propagation and cloning: 25μg/mL kanamycin, 25μg/mL chloramphenicol, 50μg/mL apramycin, 25μg/mL zeocin, 1.25μg/mL EMB, and 1.25μg/mL TCA1.

### Plasmid construction

Plasmids were constructed using isothermal assembly (ITA) or site-directed mutagenesis by KLD reaction (NEB) (**Table S2**) and transformed into *E. coli* DH5α (λ_pir_) or NEB10-beta (NEB) cloning strains by heat shock (42°C for 40 seconds) or electroporation, respectively. Primers used in plasmid construction are listed in **Table S3** and were purchased from IDT.

### Strain construction

*C. glutamicum* competent cells were prepared as previously described (32, 46, 103). Briefly, cells were sub-cultured in competency medium (BHI supplemented with 91g sorbitol, 0.1% Tween 80, 0.4g isoniazid, 25g glycine) at a 1:50 or 1:100 dilution and grown at 30°C for 3 hours. Cells were washed twice in cold 10% glycerol spinning at 3,000rpm for 10 minutes. Gene deletion was performed using allelic exchange via *sacB* counterselection using pCRD206 derivatives as described previously (104) or by recombineering as previously reported (32). Integration of pK-PIM derivatives (pSEC1 variants) was performed as described previously (101). Colony PCR was used to check pK-PIM integration, recombineering/allelic exchange insertion, and deletions as previously detailed (32).

### Flow cytometry

Cells were grown until mid-log (∼5 hours at 30°C to reach an OD_600_ ∼ 1.5-2.0) with aeration. An equivalent of 1mL at an OD_600_ = 0.3 of cells were pelleted for 5 minutes at 15,000rpm. Pellets were resuspended in 100μL of 1X PBS containing a 1:100 dilution of anti-6x His Alexa 647 (abcam) and incubated at 25°C in the dark for 1 hour. Cells were washed three times in 100μL 1X PBS spinning at 5,000rpm for 10 minutes and resuspended to final volume of 3mL in 1X PBS. Samples were analyzed on a BD LSRII flow cytometry machine using the APC optical configuration (637nm laser, 660/20 bandpass filter).

### Immunoblot analysis

The OD_600_ of stationary phase cultures was measured and an equivalent of 1mL at an OD_600_ = 2.0 was pelleted. Pellets were resuspended in 50μL of lysis buffer (per sample: 0.5μL universal nuclease (Pierce), 0.5μL 1M MgSO_4_, 1uL of protease inhibitor cocktail (Sigma-Aldrich), 10μL of 10mg/mL lysozyme, and 8μL of H_2_O). Samples were lysed at 37°C for 1.5-2 hours and then diluted 1:2 in tris-tricine sample buffer (BioRad) or Laemmli sample buffer (BioRad) supplemented with β-mercaptoethanol. PorH-His and PorH-ALFA immunoblot samples were performed on 16.5% tris-tricine gels (BioRad) using anti-His (mouse) antibody (GenScript) at a dilution of 1:2,000-1:3,000 or anti-ALFA (HRP) antibody (Nanotag Biotechnologies). NativePAGE, purification, and mAGP sacculi samples were analyzed on 4-20% TGX gels (BioRad). mAGP sacculi, purification, and loading control sample gels were stained with ReadyBlue Coomassie stain (Sigma-Aldrich), where appropriate. In all cases, anti-mouse (HRP) antibody (Rockland Biosciences) was used at a dilution of 1:3,000-1:5,000. A BioRad Chemidoc imager was used to visualize immunoblots and FIJI (v1.54n) was used to analyze images (105).

### Fluorescence microscopy for live-cell induction experiments

To visualize MOMP localization strains H1616 (Δ*porH* P*_tac_*::*porH-ALFA*) or DB42 (P*_tac_*::*protX-ALFA*) were grown overnight in BHI supplemented with chloramphenicol (3.5μg/mL) at 30°C, 200rpm. The next day, cells were diluted 1:200 and grown until OD_600_ = 0.2-0.3 at 30°C. Samples were harvested by centrifugation (2 min, 5000xg). Cells and 12.5 nM FluoTag®-X2 anti-ALFA Atto488 labeled nanobodies (NanoTag Biotechnologies) were added onto a 2% agarose pad prepared with 1/10 BHI diluted in PBS, supplemented with 3.5µg/mL chloramphenicol and 50-500µM IPTG depending on the experiment. Diluted BHI was used to reduce autofluorescence of the media while not affecting cell growth over the indicated observation time.

Cells were imaged on a Nikon Ti2 inverted widefield microscope equipped with a fully motorized stage and perfect focus system and OkoLab environmental enclosure heated to 30°C. Images were acquired using a 1.45 NA Plan Apo x100 Ph3 objective lens with Nikon type F2 immersion oil. Fluorescence was excited using the Lumencore SpectraX LED light engine using 100ms exposure, 30-50% LED power and filtered through the Lumencor 470/24x, Semrock FF01-475/543/702 dichroic mirror and Chroma ET519/26m emission filter. Images were recorded on a Hamamatsu ORCA-Flash 4.0 C11440-22C camera (sCMOS, 16 bit, 6.5µm pixel size) using Nikon Elements (v5.30.05) acquisition software. Cells were imaged for two-hour observation period with a 2.5min acquisition frame rate.

Images were analyzed and rendered for figure or movie display using FIJI (v1.54n) (105) and MicrobeJ (v5.13p) (106). Time-lapse series were drift corrected using a customized StackReg plugin (107, 108). Kymographs were generated using the ‘Relsice’ function in FIJI. Fluorescence intensity distribution across cell length was extracted from MicrobeJ using the ‘Profile’ function. To this end, cell margins were dilated by 0.3 pixel laterally and 0.4 pixels longitudinally to obtain symmetrical peaks of the intensity traces at the cell poles. Intensity traces were exported, normalized and plotted using GraphPad Prism (v10.3.1). Original images will be uploaded on Zenodo (link accessible upon publication).

### Fluorescence recovery after photobleaching (FRAP)

To assess diffusion of MOMPs strains H1616 (Δ*porH* P*_tac_*::*porH-ALFA*) or DB42 (P*_tac_*::*protX-ALFA*) were grown as indicated above. Day cultures were grown until OD_600_ = 0.4 before samples were concentrated to OD_600_ = 20 by centrifugation (2 min, 5000g). Ten microliters of concentrated cells were then spotted on a BHI-Agar plate containing 500µM IPTG, 100µM HADA and 3.5µg/mL chloramphenicol and further incubated for 2h at 30°C. Subsequently, cells were recovered by punching them out of the agar plates using the back end of a Pasteur pipette. The agar disc was vigorously vortexed during 20sec in PBS and cells were transferred to a new tube for labeling with 12.5 nM FluoTag®-X2 anti-ALFA Atto488 nanobodies and 30µM 6TMR-Tre for 15min at room temperature. Last, cells were washed once (in PBS) before immobilizing them on a 2% agarose in PBS pads.

Cells were imaged on a Nikon AX inverted laser-scanning confocal microscope equipped with a fully motorized stage and perfect focus system and a 100X 1.45 NA objective lens with Nikon type F2 immersion oil. The microscope was controlled using Nikon Elements (v5.42.06, build 1821) acquisition software with a customized JOBS workflow for photobleaching. Images were acquired in Galvano unidirectional scanning mode, with 2x line averaging and 0.6µs pixel dwell time. FRAP was performed by bleaching a diffraction limited spot (∼ 250nm sized spot) using Nikon AX lasers at 405nm, 488nm and 561nm at 100% laser power for 2 µs dwell time per pixel. We placed the location of the bleached area evenly across the cell envelope, including approximately 50% poles and 50% side walls. Fluorescence recovery was followed during 1min postbleaching using a 2sec acquisition frame rate. In total 35 cells for PorH (from two biological replicates) and 55 cells for ProtX (from three biological replicates) were analyzed.

Images were analyzed and rendered for figure or movie display with FIJI (v1.54n). FRAP data was normalized to account for overall bleaching using the equation

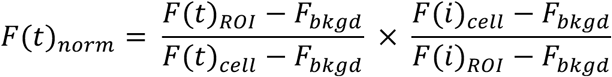

where *F*(*t*)_ROI_ and *F*(*t*)*_cell_* are the intensities of the ROI and the whole cell, respectively, at each time point *F*(*t*). Similarly, *F*(*i*)*_ROI_* and *F*(*i*)*_cell_* are the intensities of the ROI and the whole cell at the start of the experiment, while *F_bkgd_* represents fluorescence intensity outside the cell as described previously by (109). Data was visualized in GraphPad Prism (v10.3.1). Original images will be uploaded on Zenodo (link accessible upon publication).

### mAGP sacculi extraction

mAGP sacculi were isolated as previously described (55). Briefly, an equivalent OD_600_ of an overnight culture was pelleted (1.5 minutes at 15,000 x g) and resuspended in 500μL of 2% TritonX 100 in 1X PBS. Samples were sonicated in an ice-cold bath sonicator for 6 cycles of 20 seconds on/2 minutes off and then pelleted spinning for 15 minutes at 15,000 x g. Pellets were resuspended in 500μL of 2% SDS in 1X PBS and boiled at 100°C for 1 hour. Samples were divided in two, pelleted at 15,0000 x g for 15 minutes, and washed three times in 500μL each (first wash: 100% milliQ water, second wash: 80% acetone in milliQ water, third wash: 100% acetone). The resulting final pellets were resuspended in either 100uL of milliQ water or 5mg/mL lysozyme in milliQ water, with each biological sample having both a lysozyme-treated and untreated condition. Samples were incubated in a 37°C water bath for ∼16 hours. Aliquots of the mAGP sacculi were then diluted 1:2 in either tris-tricine sample buffer or Laemmli sample buffer, both supplemented with β-mercaptoethanol and analyzed by immunoblot or ReadyBlue staining. For trifluoroacetic acid (TFA) treatment of sacculi, extracted mAGP sacculi were treated with a cleavage cocktail (47.5 TFA:1.25 H_2_O:2.35 triethylsilane) for 2 hours at 25°C and neutralized to pH 7.0 with 1M NaOH. The samples were then analyzed as described above.

## ACKNOWLEDGEMENTS

The authors would like to thank members of the Bernhardt, Rudner, Walker, Vettiger, and Navarro labs for advice and helpful discussions. We would like to thank the Immunology Flow Cytometry core facility and the MicRoN imaging core at Harvard Medical School for excellent advice and maintenance of equipment. We are also grateful to Dr. Serge Pellet for maintaining the fluorescence microscopes at DMF-UNIL. This work was supported by Investigator funds from the Howard Hughes Medical Institute (to T.G.B), the Helen Hay Whitney Foundation fellowship (HHMI) (to E.M.H), the National Institutes of Health K99 AI190054 (to E.M.H), the Helen Hay Whitney Foundation fellowship (to V.M.M), the National Institutes of Health R01 148752 (to S. W.), and an Installation Credit from the University of Lausanne (to A.V.). Molecular graphics and measurements were performed using UCSF Chimera, which was developed by the Resource for Biocomputing, Visualization, and Informatics at the University of California, San Francisco with support from NIH P41-GM103311. Figures in this manuscript were made using Fiji (105), MicrobeJ (106), AlphaFold2 (36, 37), and UCSF Chimera (110).

