## Supplemental Material for "The mycomembrane proteins PorH and ProtX are inserted at polar growth zones and linked to the cell wall"

### Supplemental Materials and Methods

#### Microfluidics for pulse-chase experiments of PorH and additional cell envelope markers

To perform pulse-chase experiments cells of DB016 (*divIVa-mScarlet* $P_{tac}$::*porH-ALFA*) were grown as indicated above (Fluorescence microscopy for live-cell induction experiments). Day cultures were harvested at $OD_{600}$ = 0.1 and loaded into a CellAsic® microfluidics flow cell (Merck, B04A-03) using an ONIX pressure pump controlled by ONIX FG software. After acclimating cells for 15 minutes in 10% BHI-PBS supplemented with 3.5µg/mL chloramphenicol at 30°C, the media was switched to 10% BHI-PBS Cam3.5 supplemented with 500µM IPTG and 12.5 nM FluoTag®-X2 anti-ALFA Atto488 nanobodies (diluted 1:200 from stock) for 75 minutes. Subsequently, cells were exposed to 12.5 nM FluoTag®-X2 anti-ALFA Alexa647 nanobodies and 100µM HADA for 1 hour time-lapse with a 2.5 minute acquisition frame rate. To avoid HADA background, HADA flow was halted for 30 seconds prior to each imaging time point. Images were recorded on the same microscope as indicated above. Blue, red and far-red fluorescence were recorded using Lumencor 395/24x excitation filter, Chroma ZET405/488/561m dichroic and Chroma ET431/28m emission filter (blue, HADA), Lumencor 550/15x excitation filter, Chroma ZET405/488/561m dichroic, Chroma 49008 ET 600/50nm emission filter (red, DivIVa-mScarlet), and Lumencor 640/30x excitation filter, Semrock FF01-446/523/600/677 dichroic and Semrock FF01-684/24 emission filter (far-red, Alexa647 anti-ALFA nanobody).

Images were analyzed as above. Fluorescence intensity measurements as a function of cell length were normalized and divided into non-septating and septating cells depending on whether they display HADA incorporation at mid-cell. Data from MicrobeJ analysis was exported and visualized in GraphPad Prism (v10.3.1). This software was also used to calculate Pearson's correlation coefficient. Curve fitting for assessing fluorescence signal distribution was performed in MATLAB R2023b (MathWorks Inc.) using the *Curve Fitting* toolbox. Original images and analysis codes will be uploaded on Zenodo (link accessible upon publication).

**Fluorescence dilution assay**

To further monitor protein diffusion over longer time intervals (hours), we performed a fluorescence dilution assay. To this end we grew strains H1616 (Δ*porH* $P_{tac}$::*porH-ALFA*) or DB42 ($P_{tac}$::*protX-ALFA*) as outlined above (Fluorescence microscopy for live-cell induction experiments). ALFA-tagged protein expression was induced by supplementing day cultures with 500µM IPTG. Once the cells reached an $OD_{600}$ of 0.1, the cultures were supplemented with 100 µM HADA, 10 µM 6-TMR-tre, and a 12.5 nM dilution of Atto-488-conjugated anti-ALFA nanobody, then incubated for an additional 2 hours at 30 °C shaking at 200 rpm. The cells were then harvested (2min at 5000×g), washed twice with 1 ml BHI, and imaged on 2% agarose pads prepared with 1/10 BHI diluted in PBS. Outgrowth of cells was followed over a period of 4h with a 20min acquisition framerate using imaging conditions and system as described above (Fluorescence microscopy for live-cell induction experiments). Normalized fluorescence intensity measurements were carried out as described under "Microfluidics for pulse-chase experiments of PorH and additional cell envelope markers”. Pearson’s correlation coefficients were calculated for indicated cells using GraphPad Prism.

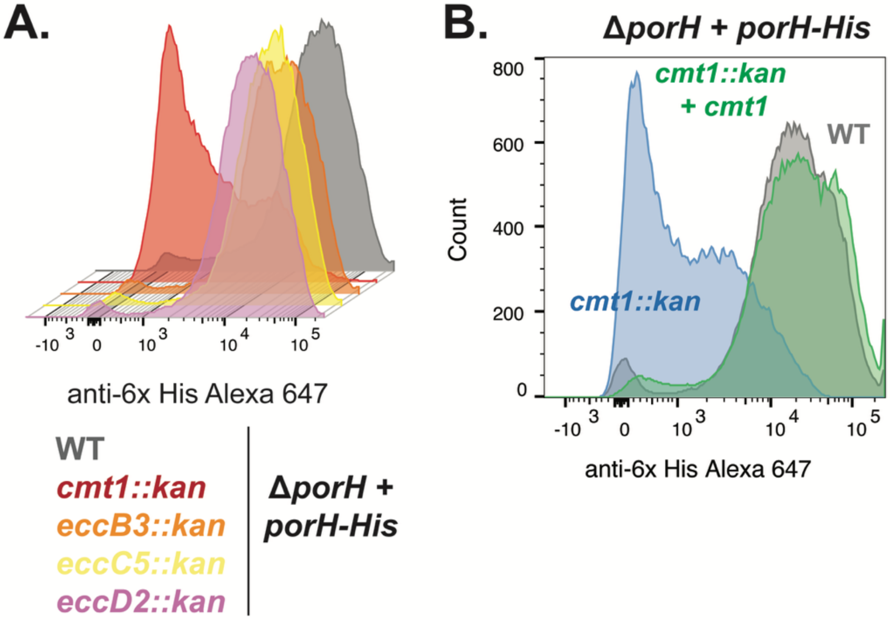


**Figure S1: Surface assembly of PorH requires Cmt1 but not the type VII secretion system.** Flow cytometry detection of PorH-His with anti-6x His antibody labeled with Alexa647. The PorH-His construct was expressed in the indicated genetic backgrounds. A representative replicate from a total of three biological replicates is displayed as a histogram. In all strains, the native copy of *porH* is deleted and PorH-His is constitutively expressed from a replicating plasmid (1, 2). **(A)** WT or cells in which genes encoding predicted type VII secretion system components were deleted. **(B)** Wild-type (WT) cells, cells lacking *cmt1*, or cells lacking *cmt1* with ectopic *cmt1* complementation. Complementation of *cmt1::kan* was performed by integrating an ectopic *cmt1* expression from a chromosomally integrated construct and inducing expression with theophylline (3, 4).

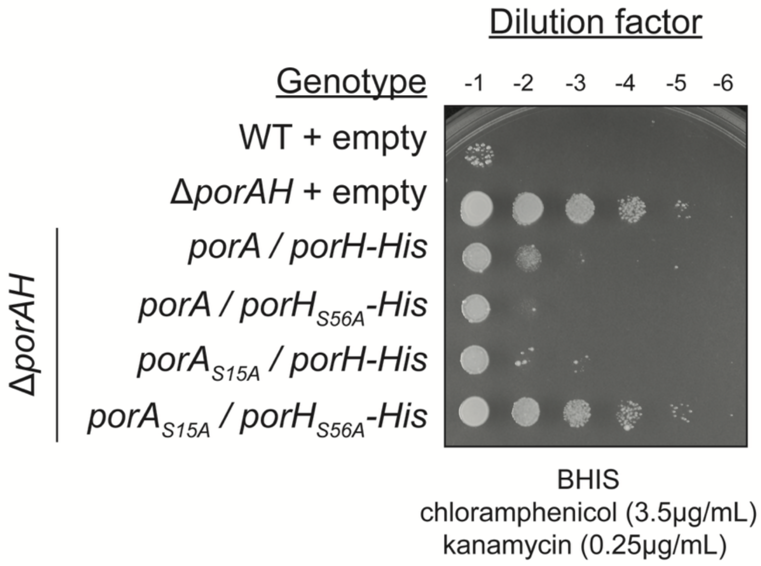


**Figure S2: *O*-mycoloylation of PorH or PorA is required for PorAH function.** Ten-fold serial dilutions of the indicated strains were spotted onto BHIS media containing 0.3μg/mL kanamycin. The *porA/porH-His* alleles were constitutively expressed from a replicating vector in a background lacking the native *porAH* locus and "+ empty" indicates that the strain carries an empty vector control. Chloramphenicol was included for plasmid maintenance. Performed in biological triplicate.

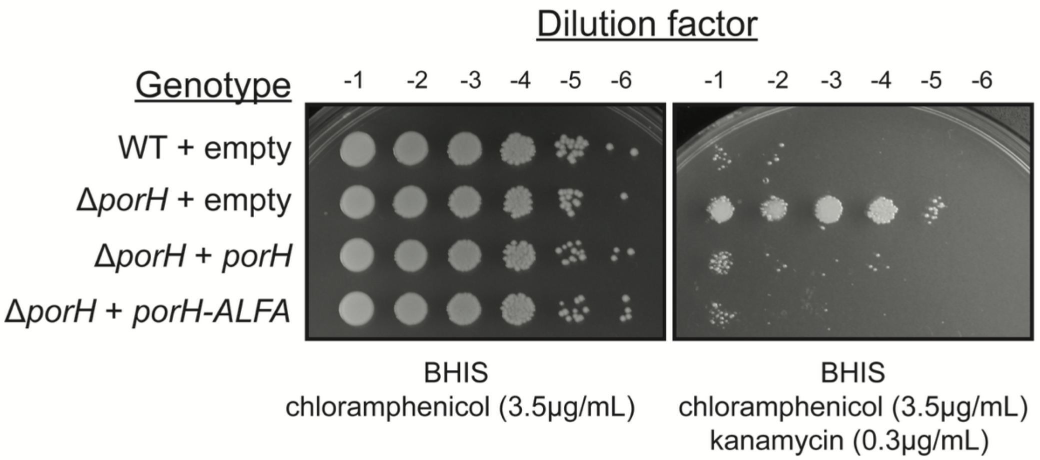


**Figure S3: Ectopic PorH-ALFA complements *porH* null kanamycin resistance.** Ten-fold serial dilutions of the indicated stains were spotted onto BHIS media or BHIS media containing 0.3μg/mL kanamycin. Strains carry either an empty vector ("+ empty") or the indicated complementation vector from which PorH or PorH-ALFA was constitutively expressed. Chloramphenicol was included for plasmid maintenance and the experiment was performed in biological triplicate.

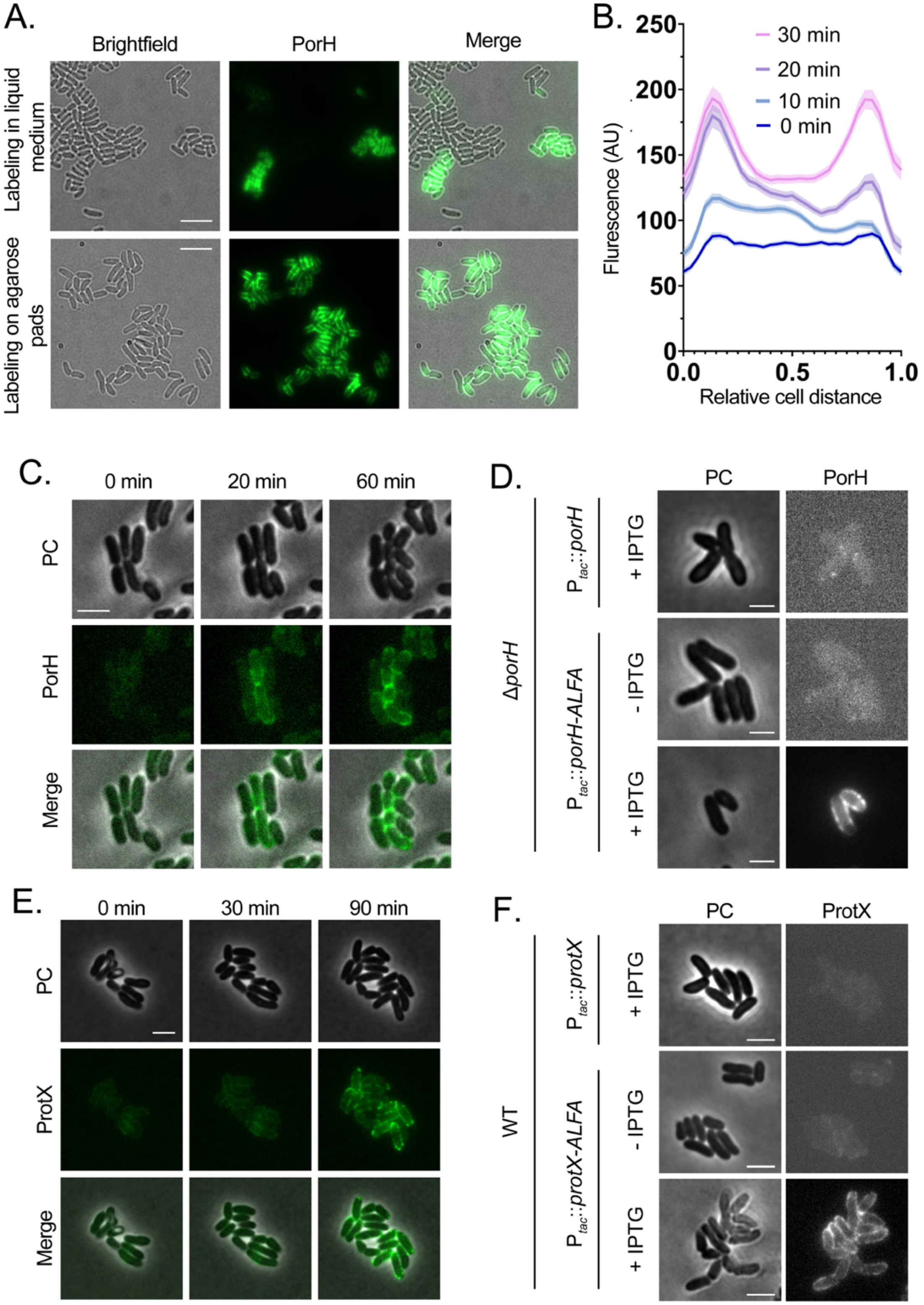

A.
Brightfield
PorH
Merge
Labeling in liquid medium
Labeling on agarose pads
B.
Flurescence (AU)
Relative cell distance
30 min
20 min
10 min
0 min
C.
0 min
20 min
60 min
PC
PorH
Merge
D.
PC
PorH
ΔporH
Ptac::porH
Ptac::porH-ALFA
+ IPTG
- IPTG
+ IPTG
E.
0 min
30 min
90 min
PC
ProtX
Merge
F.
PC
ProtX
WT
Ptac::protX
Ptac::protX-ALFA
+ IPTG
- IPTG
+ IPTG

**Figure S4: Specific detection of PorH-ALFA and ProtX-ALFA using the anti-ALFA nanobody. (A)** *Cglu* strain DB001 (Δ*porH*, $P_{tac}$::*porH-ALFA*) was induced with 500 µM IPTG for 2h in liquid (top) or on solid (bottom) BHI medium supplemented with 3.5µg/ml chloramphenicol. Cells were stained with 12.5 nM Atto488-NB and imaged by brightfield and fluorescence channel. Scale bars = 6 µm. **(B)** Quantification of Atto488-NB labeled ProtX fluorescence intensity as a function of normalized cell length at indicated time points following IPTG induction. Cells were oriented based on highest ProtX intensity (= Pole 0). Representative graph from one biological replicate (n ≥ 316 cells), shaded area indicates ± 95% confidence interval **(C)** Representative images of cells (strain H1616) expressing PorH-ALFA induced with 50 µM IPTG and stained with 12.5 nM Atto488-NB at indicated timepoints. PC = phase contrast, scale bar = 3 µm. **(D)** Cells expressing untagged PorH or PorH-ALFA with no IPTG or 500 µM IPTG induction, where indicated. Scale bars = 2 µm. **(E)** Representative images of cells (strain DB046) expressing ProtX-ALFA induced with 50 µM IPTG and stained with 12.5 nM Atto488-NB at indicated timepoints. Scale bar = 3µm. **(F)** Cells expressing untagged ProtX or ProtX-ALFA imaged with no IPTG or with 500 µM IPTG induction, where indicated. Images are representative of biological triplicates. Scale bars = 3 µm.

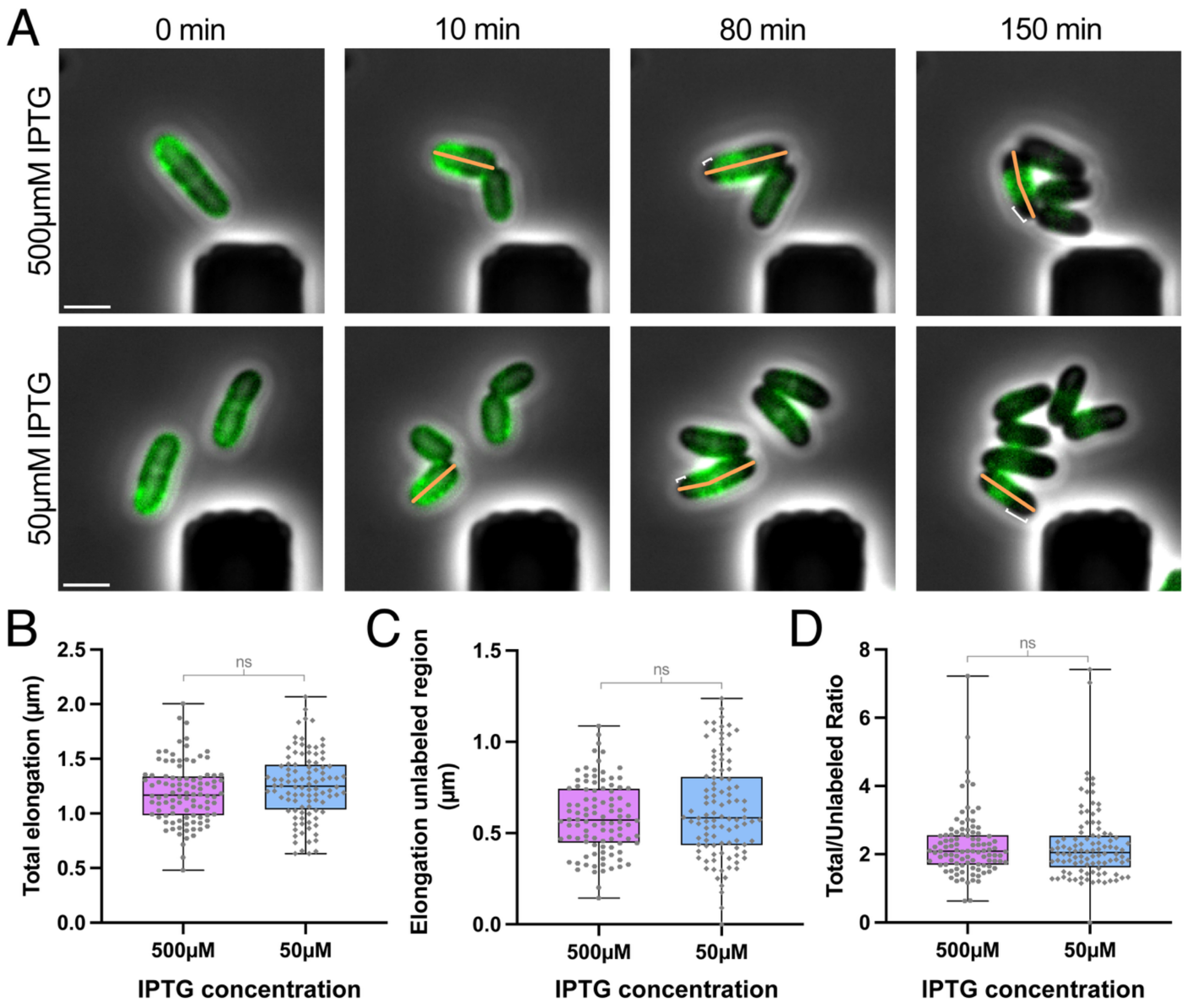


**Figure S5: Cell elongation rates and PorH incorporation are not affected by induction level**. **(A)** PorH-ALFA expression was induced with 50 or 500µM IPTG and 12.5nM FluoTag®-X2 anti-ALFA Atto488 nanobody (NB) for 5-10 minutes in a microfluidic flow cell (CellASIC) until a band of labeled PorH-ALFA was observed (denoted as '0 min'). At that point, NBs were washed out while maintaining the IPTG concentration constant. See also Materials & Methods. The orange line indicates total cell length, the white line indicates the length of the newly appearing dark poles. Scale bars are 2µm. **(B)** Quantification of the increase in cell length (orange line), as well as (**C**) newly appearing unlabeled polar material (white line) (unpaired t-test p-value = 0.10 and 0.18, respectively (ns)) and (**D**) their ratio (unpaired t-test p-value = 0.87 (ns). n = 100 cells per condition, from 2 (500µM IPTG) or 3 (50µM IPTG) biological replicates). from 2 (500µM IPTG) or 3 (50µM IPTG) biological replicates.

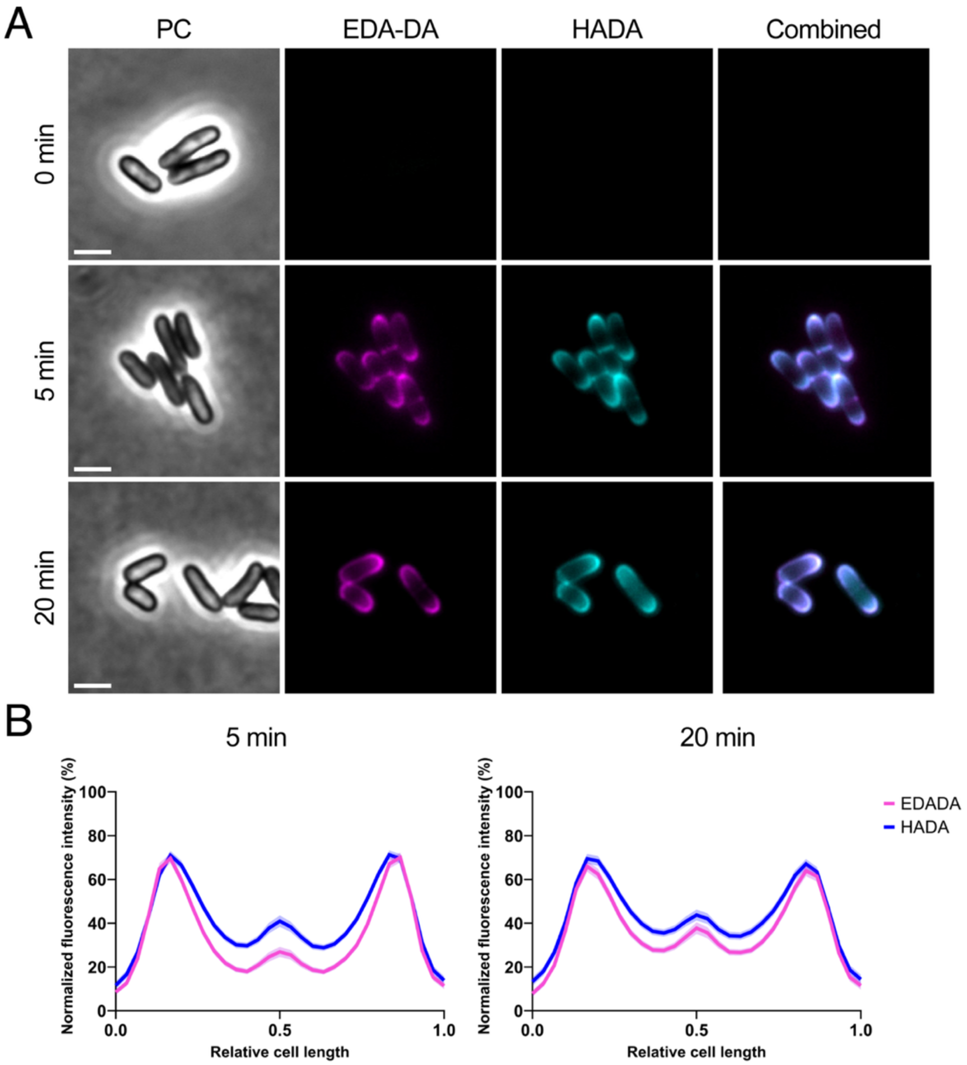


**Figure S6: Comparison of HADA and EDA-DA labelling. (A)** Representative image of cells labelled with HADA and Azide 488 labelled EDA-DA at three tested time points. **(B)** Quantification of fluorescent signal of HADA and Azide 488 labelled EDA-DA at 5 or 20 minutes. Shaded area indicates ±95% confidence interval. n = 439 (5min) or 447 (20min) from 2 biological replicates.

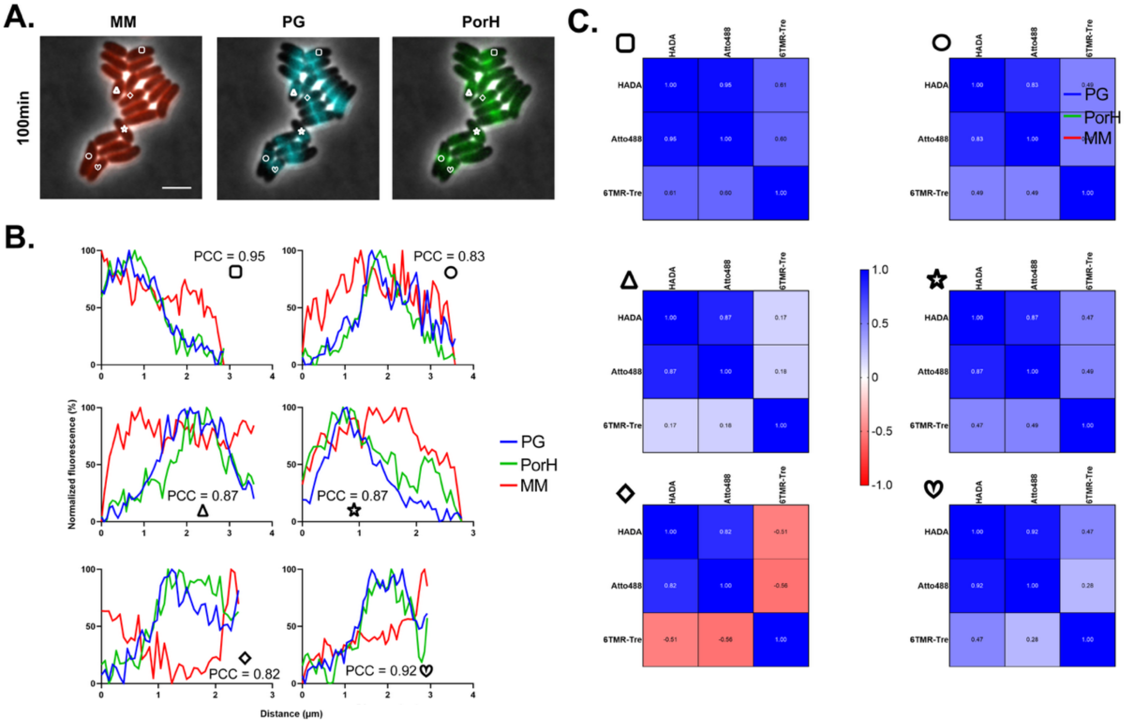


**Figure S7: PorH and PG signals strongly correlate during fluorescence dilution assay. (A)** 6TMR-tre (MM), HADA (PG), and Atto488-NB labeled PorH-ALFA (PorH) of cells identified by the indicated symbols from the fluorescence dilution assay of **Figure 4C** at the 100 minute timepoint. Scale bar is 3μm. **(B)** Normalized fluorescence intensity profiles of cells from **(A)** were plotted and Pearson correlation coefficients (PCC) determined. **(C)** Heatmap of corresponding Pearson's correlation coefficients (PCC) from line profiles in **(B)** demonstrating the stronger correlation of the PorH-ALFA and HADA signals as compared to 6TMR-tre.

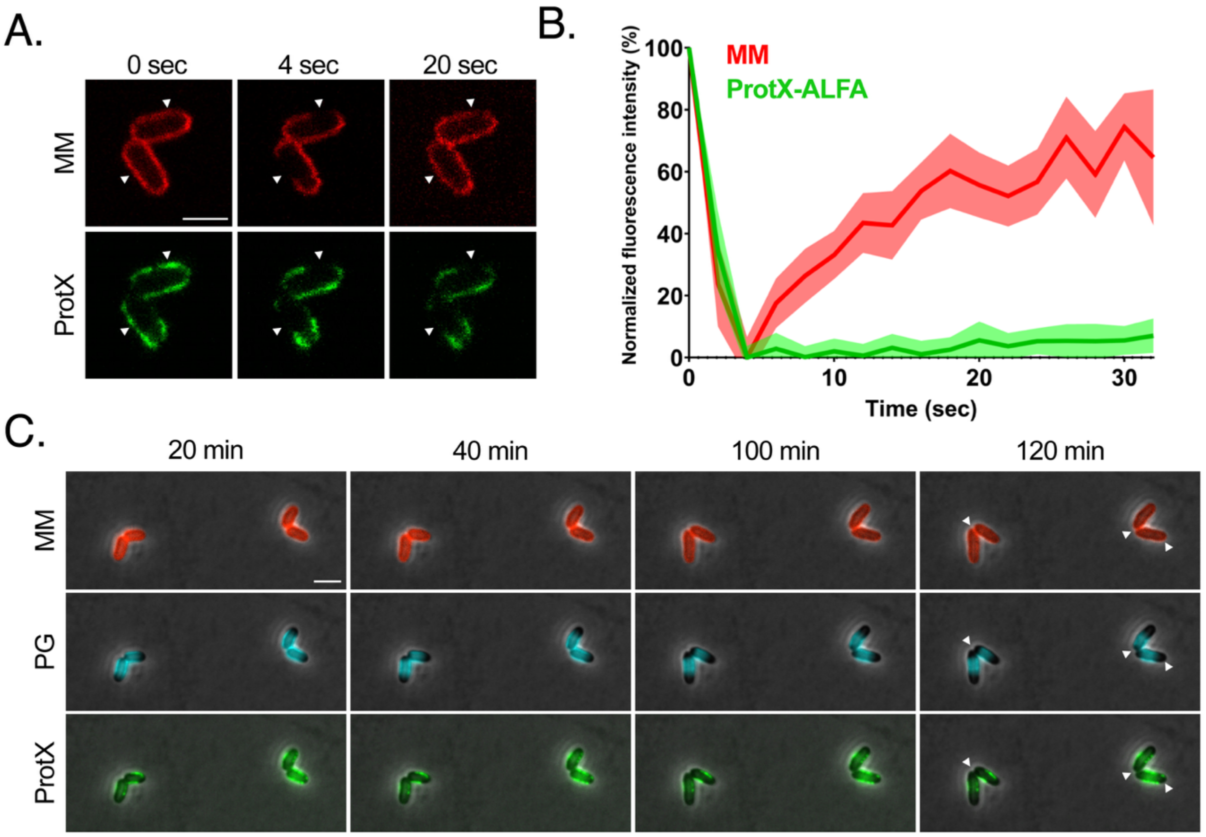


**Figure S8. ProtX remains non-diffusive upon its incorporation into the MM. (A)** Representative images of photobleached cells (strain DB42) labeled with 30 μM 6-TMR-tre (MM = mycomembrane) and 12.5 nM Atto488-NB labeling ProtX-ALFA. Arrowheads indicate bleached regions. Scale bar = 2μm. **(B)** FRAP recovery curve for MM and ProtX. Line indicates mean, shaded area indicates 95% confidence interval, n = 77 cells. **(C)** Fluorescence dilution assay for ProtX-ALFA. Cells (strain DB42) were prelabeled with same dyes as in **(A)** and imaged over 4h with 20min acquisition frame rate. Arrowheads indicate regions of new cell envelope insertion at the pole. Images are representative of three biological replicates. Scale bar = 3 μm.

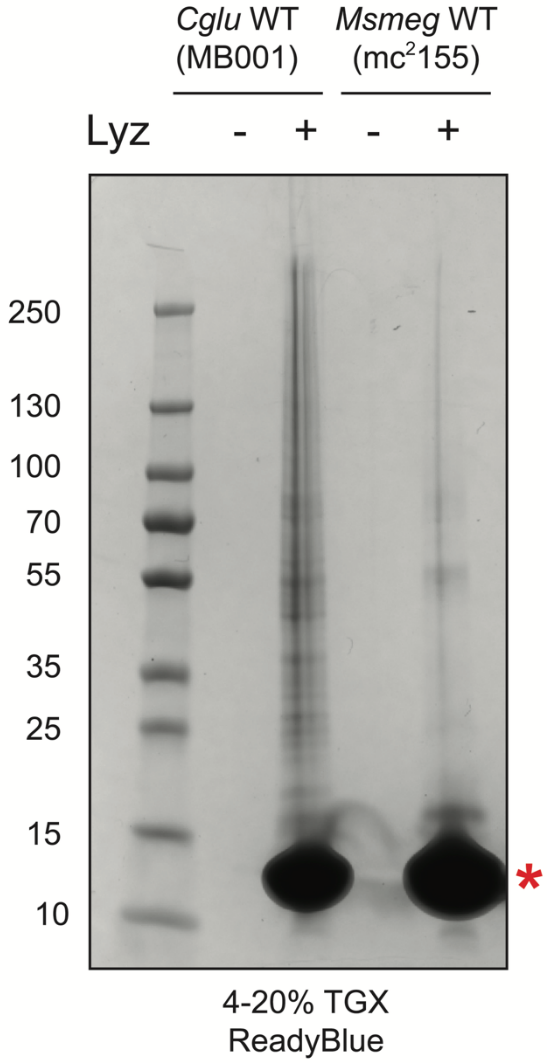


**Figure S9: Differential lysozyme susceptibility of *C. glu* and *M. smegmatis* mAGP preparations**. The mycoloyl-arabinan-peptidoglycan (mAGP) cell wall fraction was isolated from wild-type *C. glu* (MB001) or *M. smegmatis* (mc$^2$155) cells using the same extraction protocol. This fraction was left undigested or digested with 5mg/mL lysozyme ("Lyz") and analyzed by SDS-PAGE and ReadyBlue protein staining. The red asterisk highlights the band corresponding to lysozyme. This experiment was performed in biological triplicate and a representative gel image is displayed.

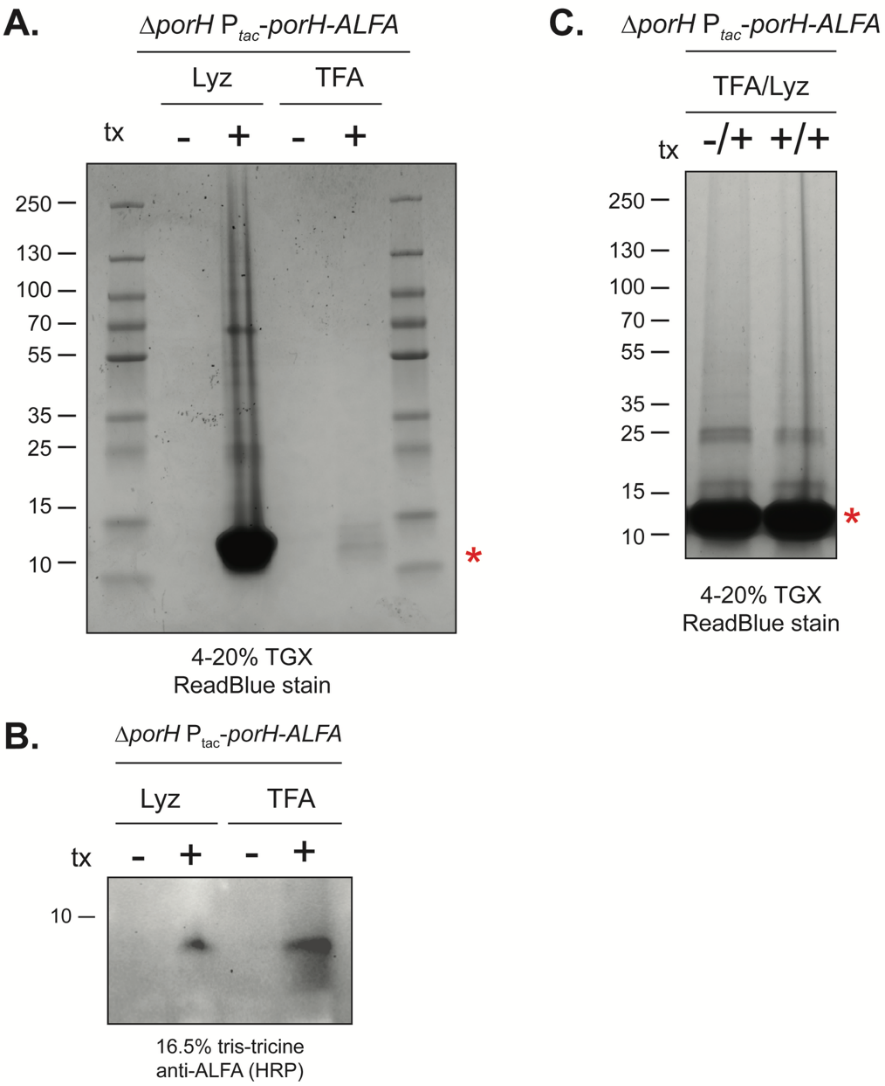


**Figure S10: PorH is released from the mAGP upon trifluoroacetic acid treatment.** The mycoloyl-arabinan-peptidoglycan (mAGP) fraction was extracted from cells expressing either untagged or ALFA-tagged PorH from an inducible promoter in the absence of the native copy of *porH*. The extracted mAGP was left untreated, digested with 5mg/mL lysozoyme overnight at 37°C, or treated with 95% trifluoroacetic acid (TFA) for 2 hours at room temperature. These samples were analyzed by, **(A)** SDS-PAGE followed by ReadyBlue staining or **(B)** immunoblot analysis. The TFA-treated sacculi were then subjected to digestion with 5mg/mL lysozyme overnight at 37°C and analyzed by SDS-PAGE and ReadyBlue staining **(C)**. The red asterisk denotes the band corresponding to lysozyme. This experiment was performed in biological triplicate.

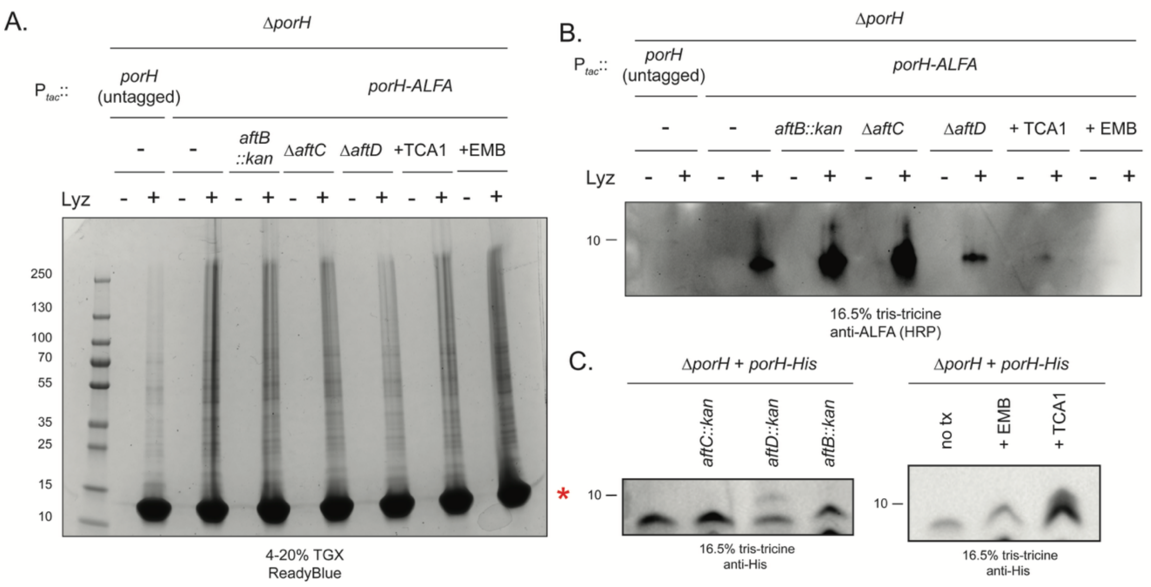


**Figure S11: Treatment with EMB or TCA1 reduces PorH association with the mAGP.** The mycoloyl-arabinan-peptidoglycan (mAGP) sacculus was extracted from cells expressing untagged or ALFA-tagged PorH expressed from an inducible promoter with 1mM IPTG induction in a genetic background lacking the native *porH* locus. Where indicated, cells were grown in the presence of 1.25μg/mL EMB or 1.25μg/mL TCA1. The genetic background of the cells is also indicated. The mAGP was left undigested or digested with 5mg/mL lysozyme and analyzed by, **(A)** SDS-PAGE followed by ReadyBlue staining or, **(B)** immunoblot analysis. **(C)** Whole cell levels of PorH in the indicated strain backgrounds were analyzed by immunoblot. The red asterisk indicates the band corresponding to lysozyme. The experiment was performed in biological triplicate and a representative replicate is shown.

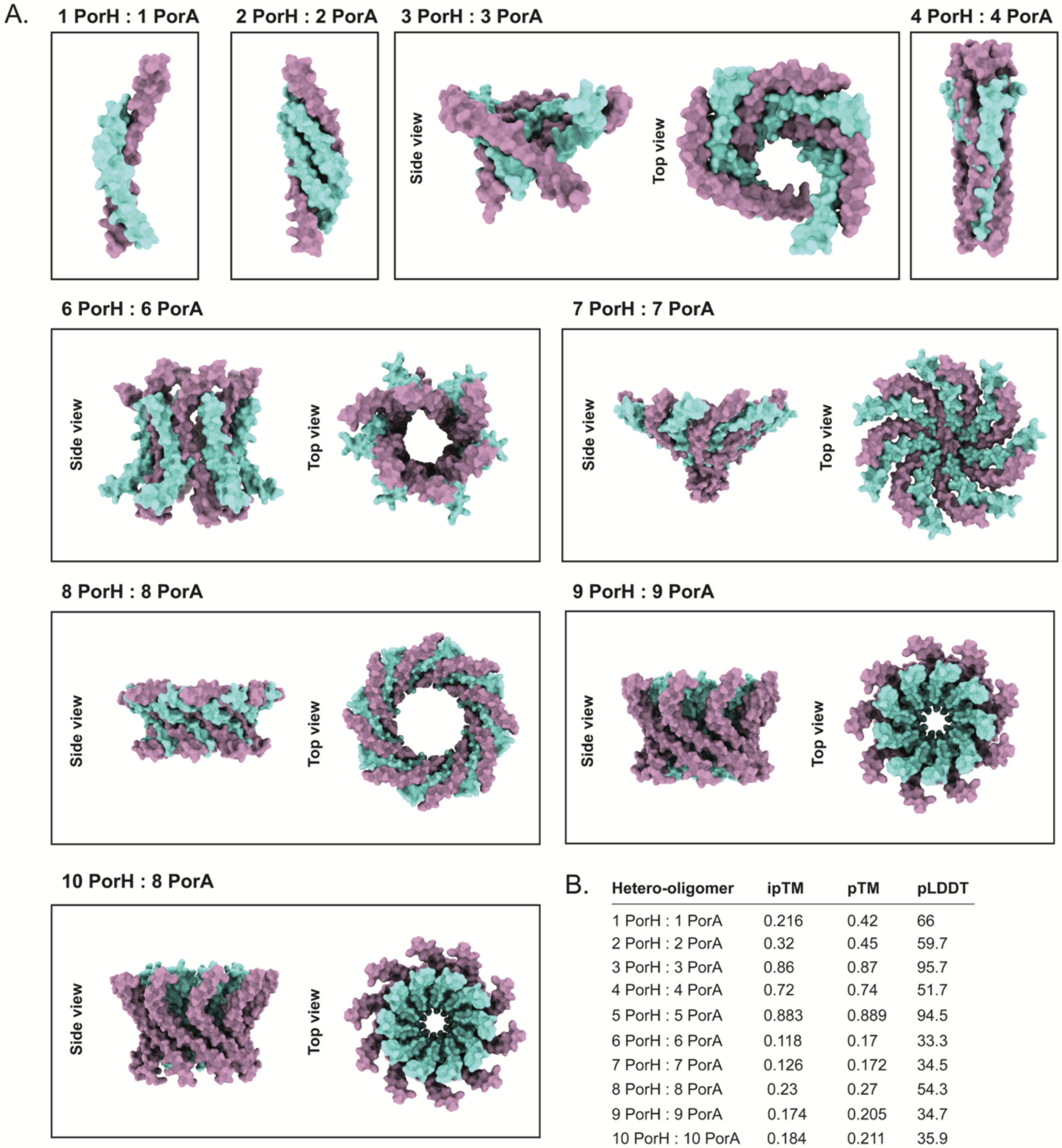


B.

| Hetero-oligomer | ipTM | pTM | pLDDT |
|---|---|---|---|
| 1 PorH : 1 PorA | 0.216 | 0.42 | 66 |
| 2 PorH : 2 PorA | 0.32 | 0.45 | 59.7 |
| 3 PorH : 3 PorA | 0.86 | 0.87 | 95.7 |
| 4 PorH : 4 PorA | 0.72 | 0.74 | 51.7 |
| 5 PorH : 5 PorA | 0.883 | 0.889 | 94.5 |
| 6 PorH : 6 PorA | 0.118 | 0.17 | 33.3 |
| 7 PorH : 7 PorA | 0.126 | 0.172 | 34.5 |
| 8 PorH : 8 PorA | 0.23 | 0.27 | 54.3 |
| 9 PorH : 9 PorA | 0.174 | 0.205 | 34.7 |
| 10 PorH : 10 PorA | 0.184 | 0.211 | 35.9 |

**Figure S12: Predicted PorAH hetero-oligomeric structures. (A)** Structural predictions of different hetero-oligomers of PorA and PorH (PorA in pink and PorH in cyan) predicted by AlphaFold2 (5). Only the Rank 1 prediction of each input is displayed (**B**) AlphaFold2 confidence measurements of the structures shown in **(A)** and Figure 6. ipTM: interface predicted Template Modeling score, pTM: predicted modeling score, pLDDT: predicted Local Distance Differenc Test.

**Supplementary Movie Legend**

**Movie S1: Live-cell imaging of PorH surface assembly.** Cells (H3922, ∆*porH* $P_{tac}$*::porH-ALFA*) were spotted on a 2% agarose pad containing 500 µM IPTG and 12.5 nM diluted Atto488-NB. PorH-ALFA surface assembly was observed over a 2 hr observation period with a 2:30 min acquisition frame rate. Three examples are shown, PC = phase contrast, scale bar = 2 µm. Movie is rendered at 10 frames per seconds.

**Movie S2: Live-cell imaging of ProtX incorporation.** Cells (DB042*,* $P_{tac}$*::protX-ALFA*) were spotted on a 2% agarose pad containing 500 µM IPTG and 12.5 nM diluted Atto488-NB. ProtX-ALFA surface assembly was observed over a 2 hr observation period following a 2:30 min acquisition frame rate. Three examples are shown, PC = phase contrast, scale bar = 2µm. Movie is rendered at 10 frames per seconds.

**Movie S3: Fluorescence recovery after photobleaching of labeled PorH-ALFA.** Cells (H3922, ∆*porH* $P_{tac}$*::porH-ALFA*) were prelabled with 30 µM 6-TMR-Tre (MM), 100 µM HADA (PG) and 12.5 nM Atto488-NB (PorH), spotted onto a 2% agarose pad, and imaged on a Nikon AX confocal microscope. Photobleaching was achieved by exposing cells to a 405nm, 488nm and 561nm laser at 100% power using 2 µs dwell time per pixel. Three examples of FRAP are shown, MM = mycomembrane, PG = peptidoglycan, scale bar = 2 µm. Movie is rendered at 7 frames per seconds.

**Movie S4: Fluorescence dilution assay of PorH.** Cells (H3922, ∆*porH* $P_{tac}$*::porH-ALFA*) were prelabled with 30 µM 6-TMR-Tre (MM), 100 µM HADA (PG) and 12.5 nM Atto488-NB (PorH), spotted onto a 2% agarose pad, and imaged over 4 hrs with a 20 min acquisition frame rate. Individual fluorescence channels are shown as merged overlays with the phase contrast channel. MM = mycomembrane, PG = peptidoglycan, scale bar = 3µm. Movie is rendered at 5 frames per seconds.

**Movie S5: Fluorescence recovery after photobleaching of ProtX.** Cells (DB042*,* $P_{tac}$*::protX-ALFA*) were prelabeled with 30 µM 6-TMR-Tre (MM) and 12.5 nMAtto488-NB (ProtX), spotted onto a 2% agarose pad, and imaged on a Nikon AX confocal microscope. Photobleaching was achieved by exposing cells to a 488nm and 561nm laser at 100% power using 2 µs dwell time per pixel. Three examples of FRAP are shown, MM = mycomembrane, scale bar = 2 µm. Movie is rendered at 6 frames per seconds.

**Movie S6: Fluorescence dilution assay of ProtX.** Cells (H3922, ∆*porH* $P_{tac}$*::protX-ALFA*) were prelabled with 30µM 6-TMR-Tre (MM), 100µM HADA (PG) and 12.5 nM Atto488-NB (ProtX), spotted onto a 2% agarose pad, and imaged over 4 hrs with a 20 min acquisition frame rate. Individual fluorescence channels are shown as merged overlays with phase contrast channel. MM = mycomembrane, PG = peptidoglycan, scale bar = 3 µm. Movie is rendered at 5 frames per seconds.

### Table S1: Strain list

| Strain number | Genotype | Strain construction | Source/reference |
|---|---|---|---|
| H60 | *C. glutamicum* MB001 (ATCC 13032 ΔCGP1 (cg1507-gp1524) ΔCGP2 (cg1746-1752) ΔCGP3 (cg1890-cg2071)) | | (6) |
| H241 | *divIVA::divIVA-mScarlet* | Replacement of native *divIVA* with *divIVA-mScarlet* fusion | (7) |
| H1111 | Δ*protX (cgp_2875)* | Deletion of *protX* (allelic exchange) | (1) |
| H1177 | MB001 pEMH25 | H60/pEMH25 | This study |
| H1241 | Δ*porH (cgp_3009)* | Deletion of *porH* (allelic exchange) | (1) |
| H3249 | Δ*porH* pEWL103 | H1241/pEWL103 | This study |
| H1248 | Δ*porH* pEMH25 | H1241/pEMH25 | (1) |
| H1249 | Δ*porH* pEMH26 | H1241/pEMH26 | (1) |
| H1448 | Δ*porH* pEMH27 | H1241/pEMH27 | (1) |
| H3369 | Δ*porH* pEMH27 pEWL103 | H1448/pEWL103 | This study |
| H3398 | Δ*porH eccB3::kan* pEMH27 | H3369 | This study |
| H3399 | Δ*porH eccC5::kan* pEMH27 | H3369 | This study |
| H3400 | Δ*porH eccD2::kan* pEMH27 | H3369 | This study |
| H4079 | Δ*porH aftC::kan* pEMH27 | H3369 | This study |
| H4070 | Δ*porH aftD::kan* pEMH27 | H3369 | This study |
| H4069 | Δ*porH aftB::kan* pEMH27 | H3369 | This study |
| H1616 | Δ*porH* pEMH145 | H1241/pEMH145 | This study |
| H5052 | Δ*porH attB2(Zeo)::EV* | H1241/pEMH691 | This study |
| H5078 | Δ*porH attB2(Zeo)::EV* pEMH27 | H5052/pEMH27 | This study |

| | | | |
|---|---|---|---|
| H5063 | Δ*porH cmt1::kan attB2(Zeo)::EV* | H3521/pEMH691 | This study |
| H5079 | Δ*porH cmt1::kan attB2(Zeo)::EV* pEMH27 | H5063/pEMH27 | This study |
| H5060 | Δ*porH cmt1::kan attB2(Zeo)::cmt1* | H3521/pEMH692 | This study |
| H5080 | Δ*porH cmt1::kan attB2(Zeo)::cmt1* pEMH27 | H5060/pEMH27 | This study |
| H3922 | Δ*porH* pEMH600 | H1241/pEMH644 | This study |
| H5597 | Δ*porH* Δ*aftC* pEMH600 | H3922 | This study |
| H5598 | Δ*porH* Δ*aftD* pEMH600 | H3922 | This study |
| H5596 | Δ*porH aftB::kan* pEMH600 | H3922 | This study |
| H1152 | Δ*porA (cgp_3008)* | Deletion of *porA* (allelic exchange) | This study |
| H1247 | Δ*porAH (cgp_3008-cgp_3009)* | Deletion of *porA/porH* (allelic exchange) | This study |
| H3542 | Δ*porAH* pEWL103 | H1247/pEWL103 | This study |
| H1493 | Δ*porAH* pEMH25 | H1247/pEMH25 | This study |
| H3019 | Δ*porAH* pEMH27 | H1247/pEMH27 | This study |
| H3291 | Δ*porAH* pEMH114 | H1247/pEMH114 | This study |
| H3292 | Δ*porAH* pEMH115 | H1247/pEMH115 | This study |
| H3293 | Δ*porAH* pEMH486 | H1247/pEMH486 | This study |
| H3294 | Δ*porAH* pEMH487 | H1247/pEMH487 | This study |
| H3521 | Δ*porH cmt1::kan* | Disruption of *cmt1* in H1241 | This study |
| H3655 | Δ*porH cmt1::kan* pEMH27 | H3521/pEMH27 | This study |
| H3763 | Δ*porAH cmt1::kan* | Disruption of *cmt1* in H1247 | This study |
| H3657 | Δ*porAH cmt1::kan* pEMH27 | H3763/pEMH27 | This study |
| H3765 | Δ*porAH cmt1::kan* pEMH114 | H3763/pEMH114 | This study |
| DB016 | *divIVA::divIVA-mScarlet* pEMH600 | H241/pEMH600 | This study |
| DB042 | MB001 pDB005 | H60/pDB005 | This study |
| DB044 | Δ*protX* pDB006 | H1111/pDB006 | This study |
| DB045 | Δ*protX* pDB005 | H1111/pDB005 | This study |

| | | | |
|---|---|---|---|
| H120 | *Mycobacterium smegmatis* $mc^2155$ | | (8) |

**Table S2: Plasmids used in this study**

| Plasmid name | Information | Source |
|---|---|---|
| pEWL89 | pCRD206($Apra^R$)::$P_{sod}$-*cre* | (1) |
| pEWL103 | pCRD206($Apra^R$)::$P_{tac}$ riboE1-SSAP/SSB | (1) |
| pCRD206 | $Kan^R$, *sacB* counterselection, temperature-sensitive origin | (9) |
| pEMH10 | pCRD206::*porA* | This study |
| pEMH16 | pCRD206::*porAH* | This study |
| pEMH25 | $P_{sod}$ empty vector ($Cam^R$, pGA1 mini replicon, constitutive expression, native 6x His deleted) | (1) |
| pEMH117 | Deletion of native 6x His to match pEMH25 | (1) |
| pEMH26 | pEMH25::*porH* | (1) |
| pEMH27 | pEMH25::*porH-His* | (1) |
| pEMH87 | pEMH25::$porH_{S56A}$-*His* | This study |
| pEMH145 | pEMH25::*porH*-ALFA | This study |
| pEMH114 | pEMH25::*porH-His/porA* | This study |
| pEMH115 | pEMH25::$porH_{S56A}$-*His/porA* | This study |
| pEMH486 | pEMH25::*porH-His/*$porA_{S15A}$ | This study |
| pEMH487 | pEMH25::$porH_{S56A}$-*His/*$porA_{S15A}$ | This study |
| pEMH120 | $P_{tac}$ empty vector (pACM246-derived), *eGFP* deleted, insertion of BamHI site in MCS | (1) |
| pEMH604 | pEMH120::*porH* | This study |
| pEMH600 | pEMH120::*porH-ALFA* | This study |
| pACM185 | $P_{sod}$ riboE1 empty vector ($Kan^R$, pK-PIM derivative, theophylline inducible) | (10) |
| pACM64 | $P_{sod}$ riboE1($Kan^R$)::*cmt1* | (10) |
| pEMH691 | $P_{sod}$ riboE1 empty vector ($Zeo^R$, pK-PIM derivative, theophylline inducible) | This study |
| pEMH692 | $P_{sod}$ riboE1($Zeo^R$)::*cmt1* | This study |
| pDB005 | pEMH120::*protX-ALFA* | This study |
| pDB006 | pEMH120::*protX* | This study |

**Table S3: Oligonucleotides used in this study**

| Primer name | Sequence (5' → 3') | Description (associated plasmid) |
|---|---|---|
| BH288 | ggataacttcgcatcctaatgacctaggtg | Forward, to make S56A mutation in *porH* by site-directed mutagenesis (pEMH87) |
| BH256 | agattctcgccggtggtg | Reverse, to make S56A mutation in *porH* by site-directed mutagenesis (pEMH87) |
| BH562 | ctgcgccgccgcctgaccgaatgacctaggtgcctggcg | Forward, to insert C-terminal ALFA tag in *porH* (pEMH145) |
| BH563 | ttcttcttccaggcggctcggggaagagaagttatccagattctcgc | Reverse, to insert C-terminal ALFA tag in *porH* (pEMH145) |
| BH421 | ctaggtgcctggcggcag | Forward, to linearize pEMH27/pEMH87 vectors (pEMH114) |
| BH422 | gtcagtggtggtggtggtg | Reverse, to linearize pEMH27/pEMH87 vectors (pEMH114) |
| BH423 | ccaccaccaccaccactgacgagaaatccgatttggctg | Forward, to amplify *porA* and *porH/porA* intergenic region from chromosome (pEMH114) |
| BH424 | tactgccgccaggcacctagttagccaagcagaccgatg | Reverse, to amplify *porA* and *porH/porA* intergenic region from chromosome (pEMH114) |
| BH257 | tgatgtccttgcaggctccggcc | Forward, to mutate residue S15A in *porA* by site-directed mutagenesis (pEMH486/pEMH487) |
| BH258 | aggtttccaaggaactcgtaaac | Reverse, to mutate residue S15A in *porA* by site-directed mutagenesis (pEMH486/pEMH487) |
| BH1570 | gcgaaaggatttttacatgatggatctttcccttctcaagg | Forward, to amplify *porH-ALFA* for insertion into pEMH120 digested with AvrII/BamHI (pEMH600) |
| BH1572 | gcgctactgccgccaggcactcattcggtcaggcggcg | Reverse, to amplify *porH-ALFA* for insertion into pEMH120 digested with AvrII/BamHI (pEMH600) |

| | | |
|---|---|---|
| BH171 | tcagaattggttaaaaaggatctagg | Forward, to linearize pK-PIM derivative vectors (pEMH691/pEMH692) |
| BH172 | cgcacagatgcgtaaggag | Reverse, to linearize pK-PIM derivative vectors (pEMH691/pEMH692) |
| BH173 | tctccttacgcatctgtgcggtacctctatctggt gccctaaac | Forward, to amplify zeocin-resistance cassette for insertion into pK-PIM derivative vectors (pEMH691/pEMH692) |
| BH174 | tcctttttaaccaattctgatcagtcctgctcctc ggc | Reverse, to amplify zeocin-resistance cassette for insertion into pK-PIM derivative vectors (pEMH691/pEMH692) |
| BH1 | agtcgacctgcaggcatg | Forward, to linearize pCRD206 (pEMH10) |
| BH2 | atccaacagggacaccag | Reverse, to linearize pCRD206 (pEMH10) |
| BH7 | gcagaataaataaatcctggtgtccc | Forward, diagnostic primer for pCRD206 derived vectors |
| BH8 | gggtaacgccagggttttcc | Reverse, diagnostic primer for pCRD206 derived vectors |
| BH95 | tcctggtgtccctgttggatacctcaattgccct cccg | Forward, to amplify upstream region of *porA* (pEMH10) |
| BH96 | gcagaccgatgttttccattttaaattctcct attaagagttgag | Reverse, to amplify upstream region of *porA* (pEMH10) |
| BH25 | aatggaaaacatcggtctgcttggctaa ttaac | Forward, to amplify downstream region of *porA* (pEMH10) |
| BH26 | tgcatgcctgcaggtcgactggcggatagcaaaa cggg | Reverse, to amplify downstream region of *porA* (pEMH10) |
| BH45 | tcctggtgtccctgttggatggcgagtacgtcg acctc | Forward, to amplify upstream region of *porAH* (pEMH16) |
| BH132 | gcagaccgataagatccatgagaaatct ccttgag | Reverse, to amplify upstream region of *porAH* (pEMH16) |
| BH133 | catggatcttatcggtctgcttggctaattaac | Reverse, to amplify downstream region of *porAH,* used with primer BH26 (pEMH16) |
| BH65 | taacatttctgcaggtcaag | Forward, diagnostic primer for *porH* |

| | | |
|---|---|---|
| BH66 | tcagcaactgcgcca | Reverse, diagnostic primer for *porH* |
| BH67 | ccgctcttcagagcatcc | Forward, diagnostic primer for *porA* |
| BH68 | ggttccatctggacgcaga | Reverse, diagnostic primer for *porA* |
| BH71 | tgcggtatttcacaccgcata | Forward, diagnostic primer for pEMH25 derivatives |
| BH250 | ccggcggatttgtcctactca | Reverse, diagnostic primer for pEMH25 derivatives |
| BH1458 | aggtcagggctaccaaccacaagttca cgagggaaagacgtatgaagctttccgtga tggtaacttcacg | Forward, recombineering primer for *cmt1* |
| BH1459 | aaaggcagccggtttcaatgttaaatct gtttctaggcctctagctcaaaagccgtca attgtctgattc | Reverse, recombineering primer for *cmt1* |
| BH81 | cagagatttttggctcgtta | Forward, diagnostic primer for *cmt1* |
| BH82 | gtgactgtcgcagcaag | Reverse, diagnostic primer for *cmt1* |
| BH1304 | cctggattaggggatgtaaagatctaaagctgggg gaatacatggctgaaggctccgtgatggtaacttc acg | Forward, recombineering primer for *eccB3* (*cgp_0661*) |
| BH1305 | caccatactccttaataggtgggcgcgagcgctgt ctcttttgctaattcactagccgtcaattgtctgattc | Reverse, recombineering primer for *eccB3* (*cgp_0661*) |
| BH1306 | atgagcaggccatcggtgt | Forward, diagnostic primer for *eccB3* |
| BH1307 | cggatcaatggcaatcactc | Reverse, diagnostic primer for *eccB3* |
| BH1308 | catatagcaccggcttaacaggccggtgctattctg ttcgcatgacttcgtccgtgatggtaacttcacg | Forward, recombineering primer for *eccC5* (*cgp_2498*) |
| BH1309 | gggcgctgagcgtctaccattgcatggtcctttagc tgaaagtgaggaaagccgtcaattgtctgattc | Reverse, recombineering primer for *eccC5* (*cgp_2498*) |
| BH1310 | tctggaacgttgattacgac | Forward, diagnostic primer for *eccC5* |
| BH1311 | tagtgaatacgtccggcgc | Reverse, diagnostic primer for *eccC5* |
| BH1312 | ggactcaccgcaggtggcgtgaaagcctagacta gatactcatgagtgcttccgtgatggtaacttcacg | Forward, recombineering primer for *eccD2* (*cgp_0830*) |
| BH1313 | tctggaattatcgtggcgttggcgcaggaactttaat tagggcggcgcttagccgtcaattgtctgattc | Reverse, recombineering primer for *eccD2* (*cgp_0830*) |
| BH1314 | cttccgcatgatcctgctt | Forward, diagnostic primer for *eccD2* |

| | | |
|---|---|---|
| BH1315 | tgctgatcggaccgttggt | Reverse, diagnostic primer for *eccD2* |
| BH1001 | gcatcggcaacgcggttgcatggccgttggccatg ttgttgatggcgcattccgtgatggtaacttcacg | Forward, recombineering primer for *aftC* (*cgp_2077*) |
| BH1002 | tttgaagtctgtcatgctgtcctctcaagatggtcgtg cgttggatcagtagccgtcaattgtctgattc | Reverse, recombineering primer for *aftC* (*cgp_2077*) |
| BH947 | tgggcccattgcatgtttatggatcta | Forward, diagnostic primer for *aftC* |
| BH948 | tgcagatctgattaaggttcttcctta | Reverse, diagnostic primer for *aftC* |
| BH975 | ttcactcggcgcatttctatgtctggattgtgctgggttt tgtggtgttttccgtgatggtaacttcacg | Forward, recombineering primer for *aftD* (*cgp_3161*) |
| BH976 | gcctagaatcgtcagaatgaactgacacccattta gcgctttggaggcctagccgtcaattgtctgattc | Reverse, recombineering primer for *aftD* (*cgp_3161*) |
| BH943 | cggtgcgggaggggtg | Forward, diagnostic primer for *aftD* |
| BH944 | gtgcgtggaagtggcgtttt | Reverse, diagnostic primer for *aftD* |
| BH1124 | ggccacatgcggtgctagcatgtggcctcatgacg tttagcccccagcgttccgtgatggtaacttcacg | Forward, recombineering primer for *aftB* (*cgp_3187*) |
| BH1125 | gggtagttatcacagcccaatgatttgcgaataagt gtttactgagagctagccgtcaattgtctgattc | Reverse, recombineering primer for *aftB* (*cgp_3187*) |
| ACM410 | cgttgacgttgaatccaccg | Forward, diagnostic primer for *aftB* |
| ACM411 | gtagtgttctgcttcggtgc | Reverse, diagnostic primer for *aftB* |
| DB007 | gtgcctggcggcagtag | Forward, to linearize pEMH120 vectors |
| DB008 | catgtaaaaaatcctttcgctagcaaattg | Reverse, to linearize pEMH120 vectors |
| DB016 | gatgcctggcagttccctac | Forward, diagnostic primer for insertions into pEMH120 |
| DB017 | cgcactcccgttctggataatg | Reverse, diagnostic primer for insertions into pEMH120 |
| DB040 | ccgagccgcctggaagaagaactgcgccgcc gcctgaccgaatgagtgcctggcggcagtag | Forward, to incorporate ALFA into linearized pEMH120 vectors |
| DB041 | gagcggataacaatttgctagcgaaaggattt tttacatgatgacctctgtattcgatatcatc | Forward, to amplify ProtX for insertion into pEMH120 linearized with DB017 and DB040 (pDB005) |
| DB042 | tcattcggtcaggcggcggcgcagttcttcttc caggcggctcggggaagaagcggtaacgatg | Reverse, to amplify ProtX-ALFA for insertion into pEMH120 linearized with DB017 and DB040 (pDB005) |

| | | |
|---|---|---|
| DB070 | tcaggtgggaccaccgcgctactgccgcca ggcacctaggaagaagcggtaacgatg | Reverse, to amplify ProtX for insertion into pEMH120 linearized with DB016 and DB017 (pDB006) |

### Supplemental Materials & Methods

#### Single-color pulse-chase experiment assessing the effect of IPTG induction levels

To assess whether different induction levels of PorH-ALFA affect rates of PorH displacement towards the sidewall, H1616 (Δ*porH* $P_{tac}$::*porH-ALFA*) cells were grown to $OD_{600}$ = 0.1 and loaded into a CellAsic® microfluidics flow cell as described above. Cells were incubated for 5-10 minutes in 10% BHI-PBS supplemented with 3.5 μg/mL chloramphenicol for plasmid maintenance, 12.5nM FluoTag®-X2 anti-ALFA Atto488 nanobodies (NBs), and either 50 or 500 μM IPTG. The medium was then switched to 10% BHI-PBS containing 3.5 μg/mL chloramphenicol and the respective IPTG concentration (50 or 500 μM) and maintained under constant flow for a 150-minute chase period at 30°C. Images were acquired at 10-minute acquisition intervals as described above.

Cells were segmented using MicrobeJ (11). Total increase in cell length ($\Delta L$) was quantified by:

$$\Delta L = Lt_{div} - Lt_0$$

Where $Lt_0$ stands for the initial cell length after the nanobody pulse and $Lt_{div}$ represents the length at the last frame before cell division (**Fig. S5B**). Similarly, the length increase of the newly incorporated unlabeled material (dark area) was measured ($L_{dark}$) (**Fig. S5C**) The ratio was calculated as (($\Delta L/L_{dark}$) (**Fig. S5D**).

**Correlating FDAA incorporation to EDA-DA probes for measuring cell wall biogenesis**

To compare HADA and EDA-DA labeling, wild-type *Cglu* MB001 (H60) cells were grown at 30°C to $OD_{600}$ = 0.3-0.4 in BHI shaking at 200rpm. Cells were then concentrated to $OD_{600}$ = 0.5 and incubated with 250 μM HADA and 250 μM EDA-DA for 0, 5, or 20 minutes before being fixed in 4% formaldehyde for 30 minutes on ice. Fluorescence labeling of EDA-DA was carried out using Click-Ti™ Protein Reaction Buffer Kit (ThermoFisher) and 5 μM of Alexa Fluor™ 488 Azide (ThermoFisher) according to manufacturer instructions. Cells were added onto a 2% agarose pad in PBS and imaged as described above. Normalized fluorescence intensity measurements were carried out as described under "Microfluidics for pulse-chase experiments of PorH and additional cell envelope markers".
